# The patient-specific mouse model with *Foxg1* frameshift mutation provides insights into the pathophysiology of FOXG1 syndrome

**DOI:** 10.1101/2025.01.21.634140

**Authors:** Shin Jeon, Jaein Park, Ji Hwan Moon, Dongjun Shin, Liwen Li, Holly O’Shea, Seon Ung Hwang, Hyo-Jong Lee, Elise Brimble, Jae W. Lee, Stewart Clark, Soo-Kyung Lee

## Abstract

Single allelic mutations in the forebrain-specific transcription factor gene *FOXG1* lead to FOXG1 syndrome (FS). To decipher the disease mechanisms of FS, which vary depending on FOXG1 mutation types, patient-specific animal models are critical. Here, we report the first patient-specific FS mouse model, *Q84Pfs* heterozygous (Q84Pfs-Het) mice, which emulates one of the most predominant FS variants. Remarkably, Q84Pfs-Het mice recapitulate various human FS phenotypes across cellular, brain structural, and behavioral levels, such as microcephaly, corpus callosum agenesis, movement disorders, repetitive behaviors, and anxiety. Q84Pfs-Het cortex showed dysregulations of genes controlling cell proliferation, neuronal projection and migration, synaptic assembly, and synaptic vesicle transport. Interestingly, the FS-causing *Q84Pfs* allele produced the N-terminal fragment of FOXG1, denoted as Q84Pfs protein, in Q84Pfs-Het mouse brains. Q84Pfs fragment forms intracellular speckles, interacts with FOXG1 full-length protein, and triggers the sequestration of FOXG1 to distinct subcellular domains. Q84Pfs protein also promotes the radial glial cell identity and suppresses neuronal migration in the cortex. Together, our study uncovered the role of the FOXG1 fragment derived from FS-causing *FOXG1* variants and identified the genes involved in FS-like cellular and behavioral phenotypes, providing essential insights into the pathophysiology of FS.

## Introduction

FOXG1 is the forkhead family transcription factor that is upregulated as the telencephalon begins to form in embryos ^1^. The elimination of the *Foxg1* gene in mice results in a striking loss of the forebrain volume ^2^, highlighting the crucial role of Foxg1 in forebrain generation.

The cortex comprises six cortical layers occupied by distinct subtypes of cortical projection neurons produced from the neural progenitors within the dorsal telencephalon ^3^. In addition to excitatory projection neurons, the cortex contains inhibitory GABAergic interneurons that are generated in the ventral telencephalon and migrate tangentially to the developing cortex ^4^. FOXG1 plays a critical role in multiple steps of cortex development. FOXG1 is vital for adequately patterning the telencephalon at early stages ^5–7^. In neural progenitor cells (NPCs), FOXG1 is required for the proliferation and maintenance of the progenitor population by blocking cell cycle exits and premature neuronal differentiation ^2,5,8,9^. In cortical neurons, FOXG1 plays a crucial role in differentiation, migration, subtype specification, and axon projection ^10–14^. The specific and complete elimination of FOXG1 in postmitotic cortical projection neurons results in inverted cortical layers ^11^, indicating that FOXG1 is needed for proper cortical lamination. Notably, deleting only a copy of the *Foxg1* gene in cortical projection neurons is sufficient to attenuate the production of upper-layer neurons, changing the composition of cortical layers ^11^. Deleting an allele of the *Foxg1* gene also leads to reduced cortical interneurons ^15^, while the complete inactivation of *Foxg1* in cortical interneurons interferes with their proper integration into the cortical layer ^16^. Therefore, FOXG1 coordinates the orderly production, migration, and integration of glutamatergic excitatory neurons and GABAergic inhibitory interneurons in the developing cortex. Importantly, these studies highlight that both copies of the *Foxg1* gene are required for building the functional cortex.

In mice, neuronal migration and layer formation are primarily completed by birth, and glial cells begin to emerge ^3^. Oligodendrocyte precursor cells (OPCs) in the cortex proliferate and differentiate into myelinating oligodendrocytes after birth, and the myelination continues to adulthood ^17^. In humans, the pathogenic variants in the *FOXG1* gene lead to the debilitating neurodevelopmental disorder collectively termed FOXG1 syndrome (FS) ^18–24^. Most FS is caused by de novo mutations in a single allele of the *FOXG1* gene. FS patients exhibit structural brain abnormalities, such as microcephaly and corpus callosum agenesis, and a delay in myelination. FS is also characterized by severe intellectual disability, hyperkinetic-dyskinetic movement disorder, irritability, and epilepsy. Many FS patients have autistic features, such as repetitive movements, poor social interaction skills, and a near absence of verbal speech and language development. Hence, FS belongs to the autism spectrum disorder (ASD). Currently, the molecular and cellular mechanisms leading to the pathology of FS remain elusive.

The critical role of FOXG1 in forebrain development and its implications in human pathogenesis raise several important questions. First, what cellular changes initiate the symptoms of FS? Second, what molecular alterations drive the pathology of FS? Third, do *FOXG1* gene variants associated with FS result in mutant forms of the FOXG1 protein, and if so, how do these mutants contribute to FS pathology? Lastly, are the processes of oligodendrocyte differentiation and maturation affected by FOXG1 mutations?

Notably, the diverse manifestations of FS are correlated with the specific types and locations of the *FOXG1* gene mutations^19,25^. This highlights the need for developing animal models that precisely replicate the genetic mutations responsible for FS and potentially produce abnormal FOXG1 protein products. Such models are crucial for understanding the pathophysiological mechanisms of the syndrome and for developing targeted treatment strategies. Existing *Foxg1-*null mouse lines^2,9,26,27^ are inadequate for this purpose, as they involve a complete deletion of the *Foxg1* gene, which does not fully capture the range of mutations seen in FS.

In this study, we report the first patient-specific FS mouse model that precisely mimics the predominant genetic variation of FS; Q84Pfs-Het mice that carry a heterozygous frameshift mutation in the *Foxg1* gene. Our research suggests that the *Q84Pfs* mutation, which corresponds to the *Q86Pfs* variant in humans, results in the expression of an N-terminal fragment of the FOXG1 protein. This fragment modifies the intracellular localization of FOXG1, promotes radial glial cell (RGC) progenitor identity, and suppresses radial neuronal migration. Remarkably, Q84Pfs-Het mice recapitulate not only FS-causing genotype but also diverse human FS symptoms, spanning from cellular anomalies to behavioral disorders such as impaired coordination and ASD-like behaviors. By studying the dysregulated genes in the cortex of Q84Pfs-Het mice, we identified the new molecular pathways involved in FS that affect neurons and oligodendrocyte lineage. In sum, our study with the patient-specific FS mouse model provides essential insights into the FOXG1-directed gene regulatory pathways and the pathophysiology of FS. Moreover, this mouse model lays the groundwork for developing targeted therapeutic approaches for FS.

## Results

### The coding region of FOXG1’s N-terminus is a mutation hotspot

The human *FOXG1* gene possesses three prominently high GC-rich regions in the coding sequences for the N-terminus of the FOXG1 protein, seven C (7C, c.250-256), seven G (7G, c.454-460), and six G (6G, c.501-506) (Fig. 1a). Correspondingly, FS-causing frameshift variants are over-represented in the N-terminal region of the FOXG1 protein prior to the DNA-binding Forkhead domain. In the FOXG1 Syndrome Research Registry, frameshift variants are the most frequent, accounting for 40.6% of participants (Fig. 1b). 81% of frameshift variants occur before the DNA-binding domain (Fig. 1c). 64.4% of frameshift cases were found in the 7C, 7G, and 6G regions, indicating that these highly G/C-repetitive regions are prone to the mutations that cause FS.

**Fig. 1.**
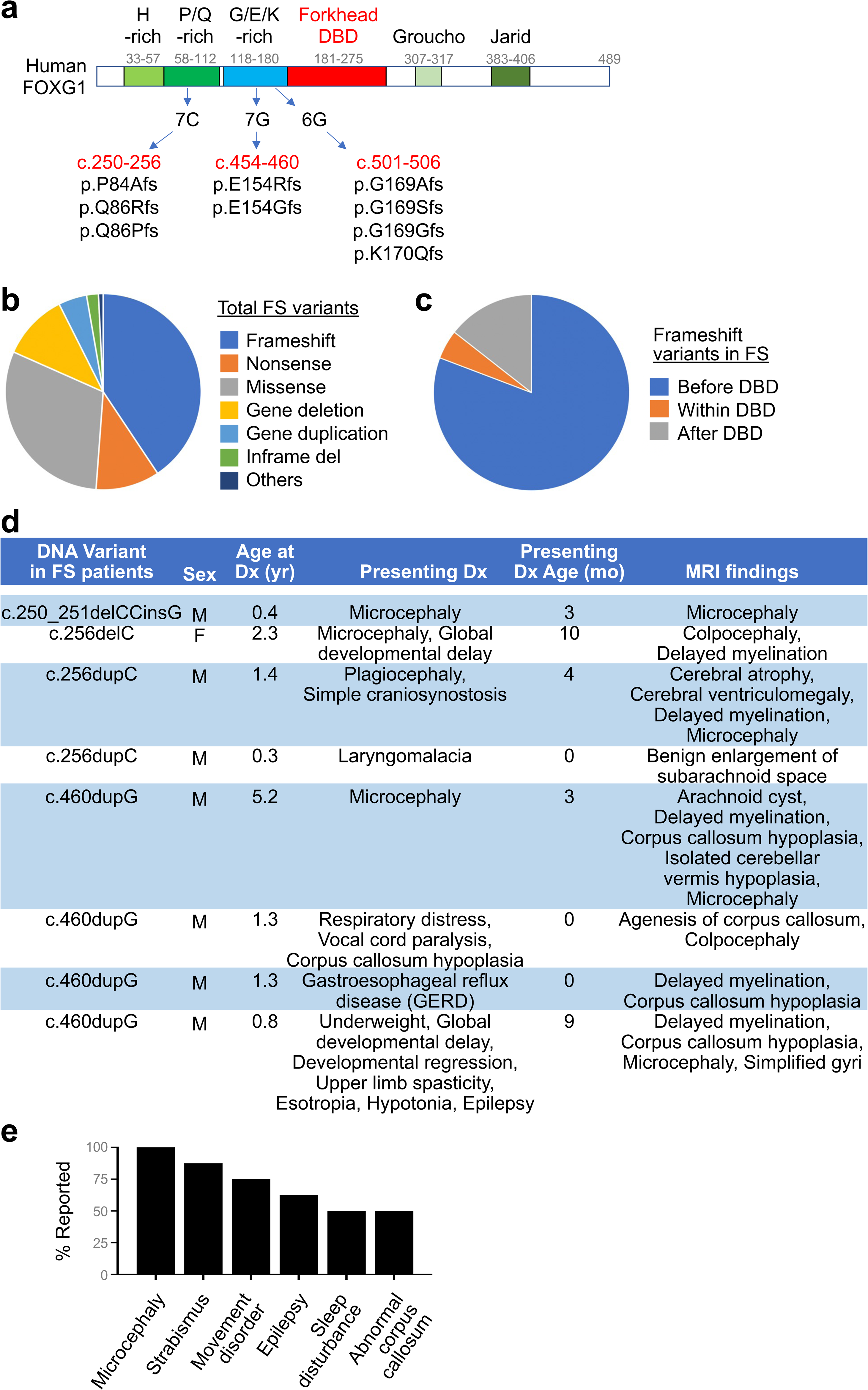
Frameshift variants causing FS reveal the three mutation hot spots in the *FOXG1* gene. **a**, Depictions of the human FOXG1 protein structure show that its N-terminal region includes domains rich in histidine (H), proline (P)/glutamine (Q), and glycine (G)/glutamic acid (E)/lysine (K). FOXG1 also contains a Forkhead DNA-binding domain (DBD) and Groucho- and Jarid-interacting domains. The three G (guanine) or C (cytosine)-repeat regions in the FOXG1-coding sequences, labeled as 7C (coding DNA reference sequences, c.250-256), 7G (c.454-460), and 6G (c.501-507), are mutation-prone areas in FS. Frameshift mutations in these regions are annotated with a “p” prefix to signify the resulting amino acid sequence changes. **b,c,** Pie charts showing the fraction of each mutation type that results in FS (**b**) and the fraction of frameshift mutations occurring before, within, or after the DBD in the FOXG1 protein (**c**). **d,** Clinical findings of eight FS patients with *FOXG1* mutations at c.250-256 and c.454-460 positions. **e,** Frequency of phenotypes among the eight FS patients in (**d**).

We reviewed the clinical data of eight individuals with FOXG1 variants at c.250-256 (7C) or c.454-460 (7G) positions (Fig. 1d,e). Over 50% of individuals with variants in these regions showed microcephaly, strabismus, movement disorder, hypotonia, epilepsy, sleep disturbance, and abnormalities of the corpus callosum agenesis (Fig. 1e). Five of eight patients also presented with delayed myelination detected via magnetic resonance imaging of the brain (Fig. 1d).

### Generation of mice carrying *Q84Pfs* allele

To understand the pathophysiology of FS, we generated FS mouse model bearing a specific FS-causing *FOXG1* variant. The knock-in mice carry c.250dupC (p.Q84Pfs*31, herein referred to as Q84Pfs), equivalent to human c.256dupC (p.Q86Pfs*35, referred to as Q86Pfs), representing one of the most prevalent variants in FS (Fig. 1a, 2a,b).

*Foxg1^Q84Pfs/Q84Pfs^* homozygous (Q84Pfs-Homo) mice exhibited a marked loss of forebrain tissue and died soon after birth (Fig. 2c, Supplementary Fig. 1a). They also showed craniofacial defects, such as a short frontonasal process, and underdeveloped eye morphology (Supplementary Fig. 1b). In contrast, *Foxg1^Q84Pfs/+^* heterozygous (Q84Pfs-Het) mice survived into adulthood.

**Fig. 2.**
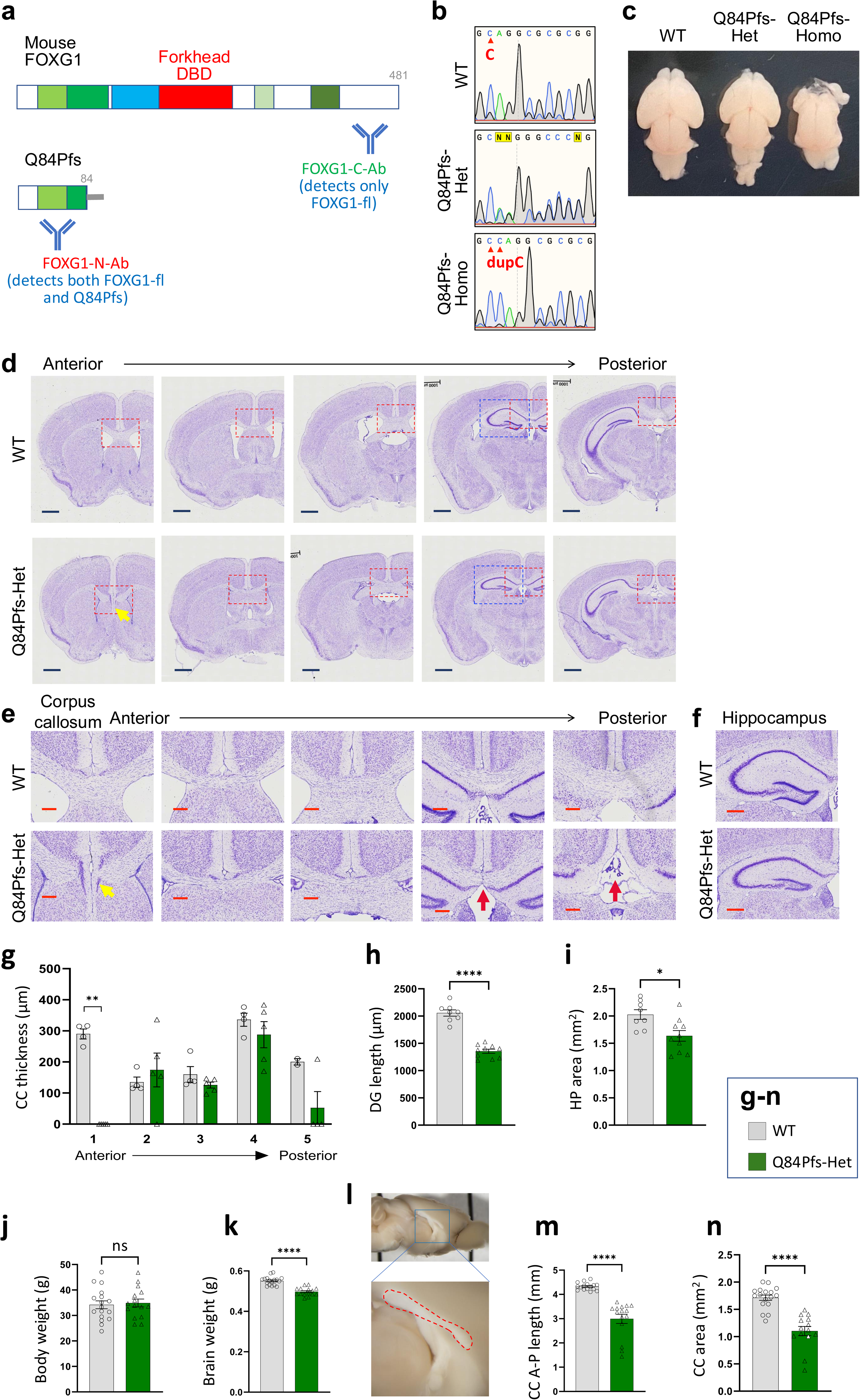
Generation of the FS animal model Q84Pfs mice and anatomical brain deficits of Q84Pfs mice. **a**, Schematics of mouse FOXG1 full-length (FOXG1-fl) protein and Q84Pfs, the truncated form of FOXG1, produced from *Foxg1* WT and Q84Pfs alleles, respectively. The antigenic regions for the two distinct FOXG1 antibodies are shown. Antibodies against the FOXG1-N-terminal region (FOXG1-N-Ab) detect both FOXG1-fl and Q84Pfs proteins, whereas antibodies against the FOXG1-C-terminal region (FOXG1-C-Ab) recognize FOXG1-fl, but not Q84Pfs. **b,** Sanger sequencing results of the *Foxg1* gene of wild-type (WT), Q84Pfs-Het, and Q84Pfs-Homo mice. An extra cytosine (C) insertion occurred after the 250th nucleotide cytosine in the Q84Pfs allele, indicated as a cytosine duplication (dupC) in the sequencing data of Q84Pfs-Homo mice. **c,** The morphology of WT, Q84Pfs-Het, and Q84Pfs-Homo brains at E18.5. **d,** The representative images for Nissl stain of the serial coronal sections of WT and Q84Pfs-Het adult brains. The anterior to posterior sections were as shown. **e,** Magnified views of the midline areas as marked as red dotted rectangles in (**d**). Yellow and red arrows indicate the septum and corpus callosum deficits, respectively. **f,** Magnified views of the hippocampus as marked as blue dotted rectangles in (**d**). **g**, Quantification of the corpus callosum thickness in the serial coronal sections of WT and Q84Pfs-Het adult brains from the anterior, corresponding to the area of the first column of (**e**), to the posterior, corresponding to the area of the first column of (**e**). **h,i**, Quantification of the length of the dentate gyrus (DG) and the size of the hippocampus (HP) areas in the serial coronal sections of WT and Q84Pfs-Het adult brains. (**g-i**, n=4 for WT, 5 for Q84Pfs-Het mice). Scale bars, 1mm (**d**) or 200μm (**e,f**). **j-n**, Quantification of body weight (**j**), brain weight (**k**), anterior-posterior length (**m**), or the total area (**n**) of the corpus callosum in the sagittal section of the brains, as depicted in (**l**) in WT and Q84Pfs-Het adult mice. (**j,k,m,n**, n=7 males, 9 females for WT, 8 males and 7 females for Q84Pfs-Het mice). The error bars (**g-k,m,n**) represent the standard deviation of the mean. \**p*<0.05, \*\**p*<0.01, \*\*\*\**p*<0.0001 and ns, non-significant in Mann-Whitney test (**g**,**m**,**n**) or unpaired two-tailed t test (**h**-**k**). Only the representative images are shown.

### Structural brain deficits in Q84Pfs-Het mice

To examine the anatomy of adult brains, we performed the Nissl staining on the coronal sections of forebrains. Compared to wild-type (WT) littermate controls, adult Q84Pfs-Het mice showed smaller brains, longitudinally shortened corpus callosum, and septum defects (Fig. 2d,e,g). Moreover, Q84Pfs-Het mice showed a significantly reduced size of the hippocampus and the malformation of the dentate gyrus (Fig. 2d,f,h,i). Consistently, adult Q84Pfs-Het mice exhibited a reduced brain weight despite no significant difference in body weights (Fig. 2j,k, Supplementary Fig. 2a,b). Interestingly, the quantification of a series of coronal sections indicates that the corpus callosum is markedly shorter in the anterior-to-posterior axis in the cross-sections of Q84Pfs-Het brains than those of control brains (Fig. 2d-g). The analysis of longitudinal length of the corpus callosum in the sagittal brain sections also revealed that Q84Pfs-Het brains show smaller and shorter corpus callosum than WT brains (Fig. 2l-n, Supplementary Fig. 2c,d). Together, Q84Pfs-Het mice recapitulate the brain structural deficits in human FS with N-terminal frameshift variants.

### The expression and function of Q84Pfs fragment

Next, we asked whether Q84Pfs protein, a truncated form of FOXG1 (Fig. 2a), is present in Q84Pfs-Het cortex and, if so, whether Q84Pfs protein plays a role in cells. To this end, we performed the immunostaining analyses in E16 Q84Pfs-Het cortex with two distinct FOXG1 antibodies, which recognize FOXG1-N-terminal region (FOXG1-N-Ab) and FOXG1-C-terminal region (FOXG1-C-Ab) (Fig. 2a) and measured the relative fluorescence intensity. FOXG1-C Ab, which detects only FOXG1 full-length (FOXG1-fl), but not Q84Pfs protein, displayed a significantly reduced fluorescence signal in Q84Pfs-Het brains compared to WT brains (Fig. 3a,b), indicating that FOXG1-fl protein levels were decreased in Q84Pfs-Het mice as expected given the heterozygous condition of the *Foxg1* gene. In contrast, the fluorescence signals of FOXG1-N-Ab, which recognizes both FOXG1-fl and Q84Pfs proteins, were slightly increased in Q84Pfs-Het brains relative to WT brains (Fig. 3a,b), suggesting the combined amount of FOXG1-fl and Q84Pfs proteins in Q84Pfs-Het cells are similar to total FOXG1-fl levels in WT cells. Intriguingly, FOXG1-N-Ab showed prominent speckles in Q84Pfs-Het neurons (Fig. 3a), suggesting that Q84Pfs protein drives intracellular speckle formation. Together, our results indicate that Q84Pfs fragment protein is expressed in Q84Pfs-Het brains.

**Fig. 3.**
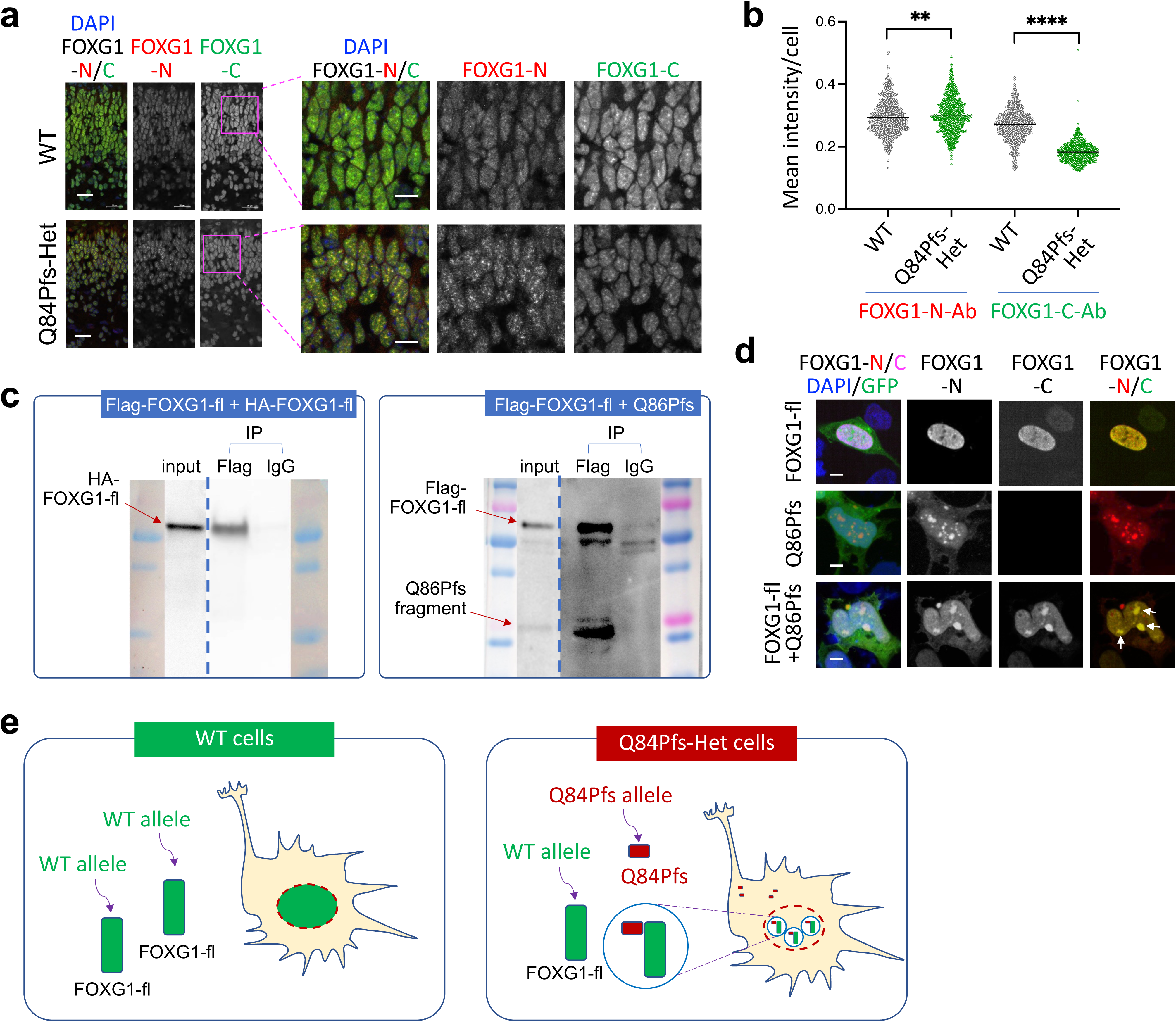
Q84Pfs fragment (Q86Pfs for human) is expressed in Q84Pfs-Het mouse cortex and interacts with FOXG1-fl. **a,b,** The immunostaining analyses of E16 Q84Pfs-Het and WT cortices with FOXG1-N-Ab and FOXG1-C-Ab. The fluorescence intensity in individual cells was quantified in (**b**). The signal intensity with FOXG1-N-Ab was comparable between Q84Pfs-Het and WT, but that with FOXG1-C-Ab was significantly lower in Q84Pfs-Het than WT. Scale bars, 25μm (a set of images on the left) or 10μm (a set of magnified images on the right). **c**, CoIP assays in HEK293 cells transfected with Flag-tagged FOXG1-fl and HA-tagged FOXG1-fl (the left set) or with FOXG1-fl and Flag-tagged Q86Pfs (the right set). In the coIP set on the left, the association between Flag-FOXG1-fl and HA-FOXG1-fl was tested by immunoprecipitation with Flag antibody, followed by western blot with HA antibody. In the coIP set on the right, the association between FOXG1-fl and Flag-Q84Pfs was monitored by immunoprecipitation with Flag antibody, followed by western blot with FOXG1-N-Ab that detects both FOXG1-fl and Q84Pfs and distinguishes the two proteins by the protein size differences. CoIP data show that FOXG1-fl interacts with each other and with Q86Pfs fragment. **d**, The immunostaining analyses in HEK293 cells transfected with FOXG1-fl alone (top panel), Q86Pfs alone (middle panel), and both FOXG1-fl and Q86Pfs (bottom panel). GFP labels the transfected cells as all plasmids have ires-GFP sequences. The subcellular distribution of FOXG1-fl and Q86Pfs proteins was assessed using FOXG1-N-Ab and FOXG1-C-Ab. FOXG1-fl protein, detected by both antibodies, is localized in nuclei, whereas Q86Pfs, recognized by only FOXG1-N-Ab but not by FOXG1-C-Ab, shows prominent speckles in nuclei and cytosol. When co-expressed, FOXG1-fl is localized in Q86Pfs^+^ nuclear speckles (white arrows). Scale bars, 5μm. **e**, The model. In Q84Pfs-Het neurons, Q84Pfs proteins are co-expressed and form speckle-like patterns with FOXG1-fl. In WT neurons, FOXG1 is primarily expressed in nuclei. The error bars (**b**) represent the standard deviation of the mean. \*\*\*\**p*<0.0001 and ns, non-significant in Mann-Whitney test. Only the representative images are shown.

To test whether human Q86Pfs (equivalent to mouse Q84Pfs) associates with FOXG1-fl, we co-expressed the two proteins in HEK293 cells and performed co-immunoprecipitation (coIP) assays. FOXG1-fl proteins were associated with each other (Fig. 3c), suggesting a dimer or multimer formation of FOXG1. Interestingly, Q86Pfs protein was co-precipitated with FOXG1-fl (Fig. 3c), indicating an interaction between Q86Pfs and FOXG1-fl in cells. Notably, in HEK293 cells, Q86Pfs protein displayed a remarkable speckle-like pattern in both nuclei and cytosol, while FOXG1-fl was expressed mainly in nuclei (Fig. 3d). When Q86fs and FOXG1-fl were co-expressed, FOXG1-N-Ab detecting both Q86Pfs and FOXG1-fl showed nuclear and cytosolic speckles. Interestingly, FOXG1-C Ab recognizing only FOXG1-fl but not Q86Pfs also displayed the overlapping speckle-like signals with FOXG1-N Ab for nuclear, but not cytosolic, speckles (Fig. 3d), indicating that FOXG1-fl is recruited to Q86Pfs-containing nuclear speckles. Together, our findings suggest that Q86Pfs fragment forms intracellular speckles and can trigger the sequestration of FOXG1-fl protein to distinct subcellular regions via associating with FOXG1-fl (Fig. 3e).

### Q86Pfs fragment suppresses neuronal differentiation and migration

To test if Q86Pfs can alter the cortical neuronal development *in vivo*, we expressed Q86Pfs protein in the developing cortex using the *in utero* electroporation of Q86Pfs-ires-GFP plasmid at E15.5 and monitored the location of GFP^+^ cells in different cellular zones of the cortex three days post-electroporation (Fig. 4a). The electroporation of the GFP plasmid was used as a control. In this experimental scheme, the location of GFP^+^ cells in the embryonic cortex reflects the status of cell differentiation ^28^. Cells in the ventricular and subventricular zones (VZ/SVZ) are NPCs, whereas those in the intermediate zone (IZ) are newly differentiated neurons that are migrating tangentially toward the cortical plate (CP). Cells in the subplate (SP) and CP represent more mature cortical neurons radially migrating to or arriving in the final location within the cortex. Interestingly, Q86Pfs protein formed speckles in cells in the developing cortex (Fig. 4a), similar to HEK293 cells. Remarkably, fewer GFP^+^ neurons entered the CP and more GFP^+^ cells were retained in the IZ in Q86Pfs-electrporated brains than in GFP alone-expressing control brains (Fig. 4a,b), indicating that Q86Pfs suppresses neuronal migration in the developing cortex.

**Fig. 4.**
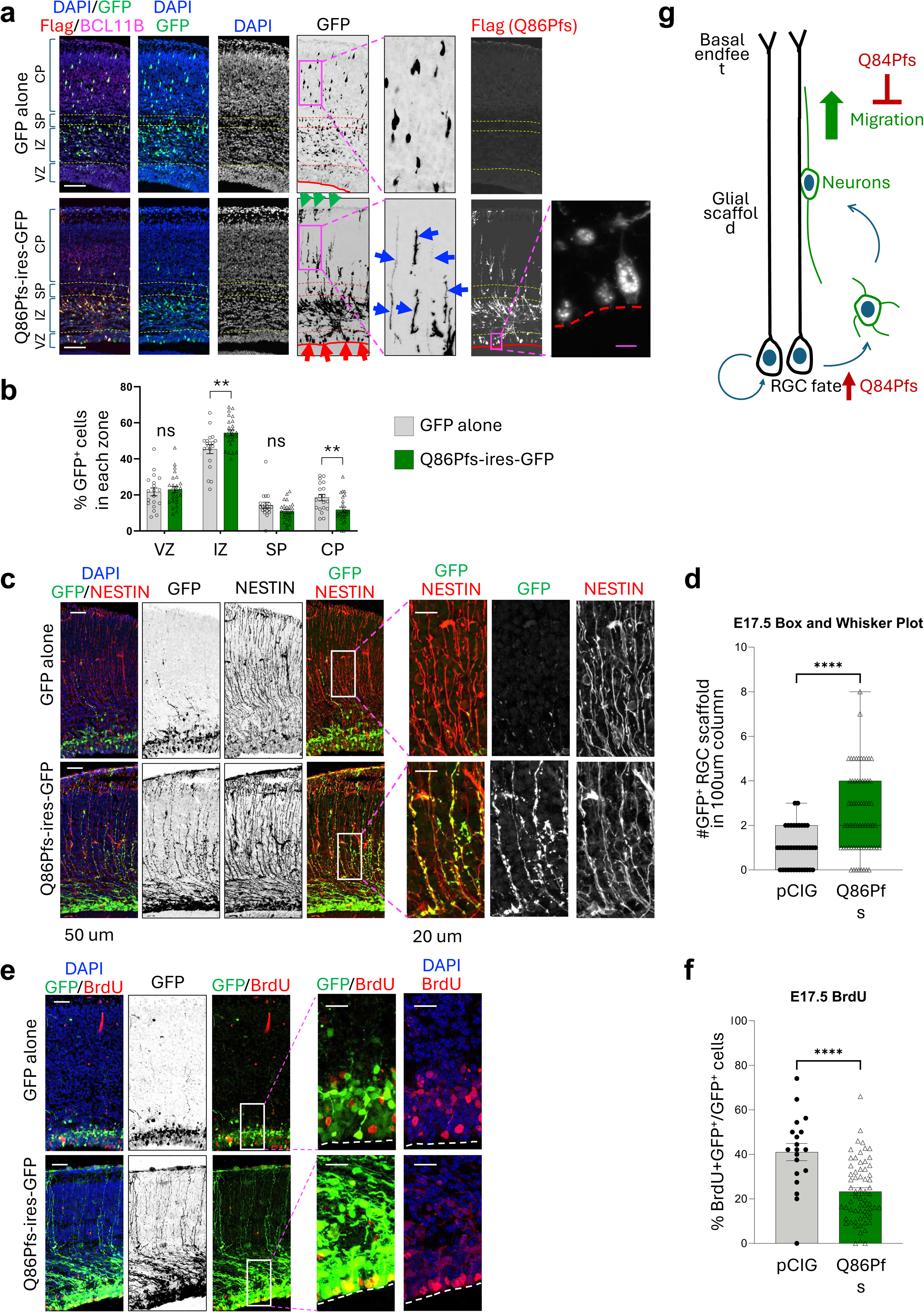
Q86Pfs fragment inhibits neuronal migration, while promoting self-renewing RGC fate. The in vivo activity of Q86Pfs was monitored in mouse brains electroporated with GFP alone or Q86Pfs-ires-GFP constructs. In utero electroporation was performed at E15.5, and the brains were harvested at E18.5 (**a**,**b**) or E17.5 (**c**-**f**). GFP labels the electroporated cells that express Q86Pfs (Q86Pfs-ires-GFP) or GFP alone. **a**,**b**, Analysis of neuronal migration of GFP^+^ cells. The proportion of GFP^+^ cells was significantly increased in the IZ and decreased in the CP in Q86Pfs-expressing brains compared to GFP-alone-expressing brains, indicating that Q86Pfs suppresses neuronal migration. BCL11B marks deep-layer neurons and was used to demarcate the CP. The immunostaining with Flag antibody shows the subcellular localization pattern of Flag-tagged Q86Pfs protein. The magnified view of the Flag staining depicts the speckle-like distribution of Q84Pfs protein. Red arrow, GFP^+^ RGC cell bodies at the ventricular surface; blue arrow, GFP^+^ RGC’s glial processes; green arrow, GFP^+^ RGC’s basal endfeet at the pial surface; red line, ventricular surface. VZ, ventricular zone; IZ, intermediate zone; SP, subplate; CP, cortical plate. **b**, Quantification of the percentage of GFP^+^ cells in each area. n=5 mice/condition. **c**,**d**, Analysis of RGC fate using the RGC marker NESTIN. The number of GFP^+^ RGC’s glial scaffold is significantly increased by Q86Pfs expression. **e,f**, Analysis of cell cycle S-phase progression using BrdU labeling. n=3 for GFP alone and n=7 for Q86Pfs-ires-GFP. The proportion of BrdU^+^ cells among GFP^+^ electroporated cells was significantly reduced by Q86Pfs expression. Scale bars, 100μm (low magnification images in a), 10μm (magnified view of Flag immunostaining, a), 50μm (low magnification images in c,e), 20μm (magnified images in c,e). The error bars (**b,d,f**) represent the standard deviation of the mean. \*\**p*<0.01, \*\*\*\**p*<0.0001 and ns, non-significant in Mann-Whitney test. Only the representative images are shown. **g**, Model for Q84Pfs actions to block neuronal migration and promote RGC fate.

We considered the possibility that Q86Pfs regulates NPC identities in addition to neuronal migration. The VZ/SVZ contains two types of NPCs: RGCs and intermediate progenitors (IPCs)^29^. RGCs extend long and thin processes to the pial surface of the cortex, forming the glial scaffold that guides the migration of neurons to the CP, while their cell bodies are located in the VZ. IPCs detach from the ventricular surface, migrate to establish the SVZ, and remove the radial glial processes. Interestingly, strong GFP expression was detected in RGC cell bodies at the ventricular surface and RGC’s glial processes and basal endfeet at the pial surface in the Q86Pfs-electroprated cortex, but not in GFP alone-expressing cortex (Fig. 4a). These results suggest that many Q84Pfs-expressing cells in the VZ maintain the RGC identity. To study the fate of GFP^+^ progenitors before they migrate from the VZ/SVZ, we performed *in utero* electroporation with a Q86Pfs-ires-GFP plasmid at E15.5 and collected the brains two days later. To track cell cycle S-phase progression, we treated the embryos with BrdU, a thymidine analog, for 2 hours before harvesting the brains. In these brains, most GFP^+^ cell bodies resided in the VZ/SVZ in both Q86Pfs-expressing and control brains (Fig. 4c). Notably, GFP^+^ glial scaffold processes, confirmed by NESTIN expression, were significantly increased in brains expressing Q86Pfs compared to GFP-only control brains (Fig. 4c,d), indicating that Q86Pfs-expressing cells are more likely to retain their RGC identity than control cells. BrdU incorporation was decreased in Q86Pfs-expressing cells compared to control cells (Fig. 4e,f). As self-expanding progenitors have a longer S-phase for DNA quality control than NPCs committed to neuronal differentiation^30^, our data indicate that Q86Pfs promotes self-renewal over neuronal differentiation in NPCs.

Together, our findings suggest that Q86Pfs protein (Q84Pfs in mice) promotes self-expanding RGC fate, inhibits neuronal differentiation, and suppresses radial migration of cortical projection neurons into the CP (Fig. 4g).

### RNA-seq analysis identified the dysregulated gene sets in Q84Pfs-Het cortex

To identify the phenotypes of Q84Pfs-Het cortex at the cellular and molecular levels, we performed RNA-seq in cortices of Q84Pfs-Het and littermate WT control mice. As neurogenesis is largely completed and the gliogenesis begins at P1 in the mouse cortex, we chose P1 cortices for RNA-seq analysis. 222 genes were significantly changed in Q84Pfs-Het cortex relative to WT cortex, representing differentially expressed genes (DEGs) (Fig. 5a, Supplementary Dataset). Interestingly, genes expressed in the cortical upper-layer projection neurons were downregulated, whereas deep-layer projection neuronal genes were upregulated (Fig. 5a,b), suggesting that the specification of cortical projection neuronal types was dysregulated in Q84Pfs-Het cortex. Many genes involved in GABAergic interneuron development were also downregulated in Q84Pfs-Het cortex (Fig. 5b), indicating defects in the development of cortical interneurons.

**Fig. 5.**
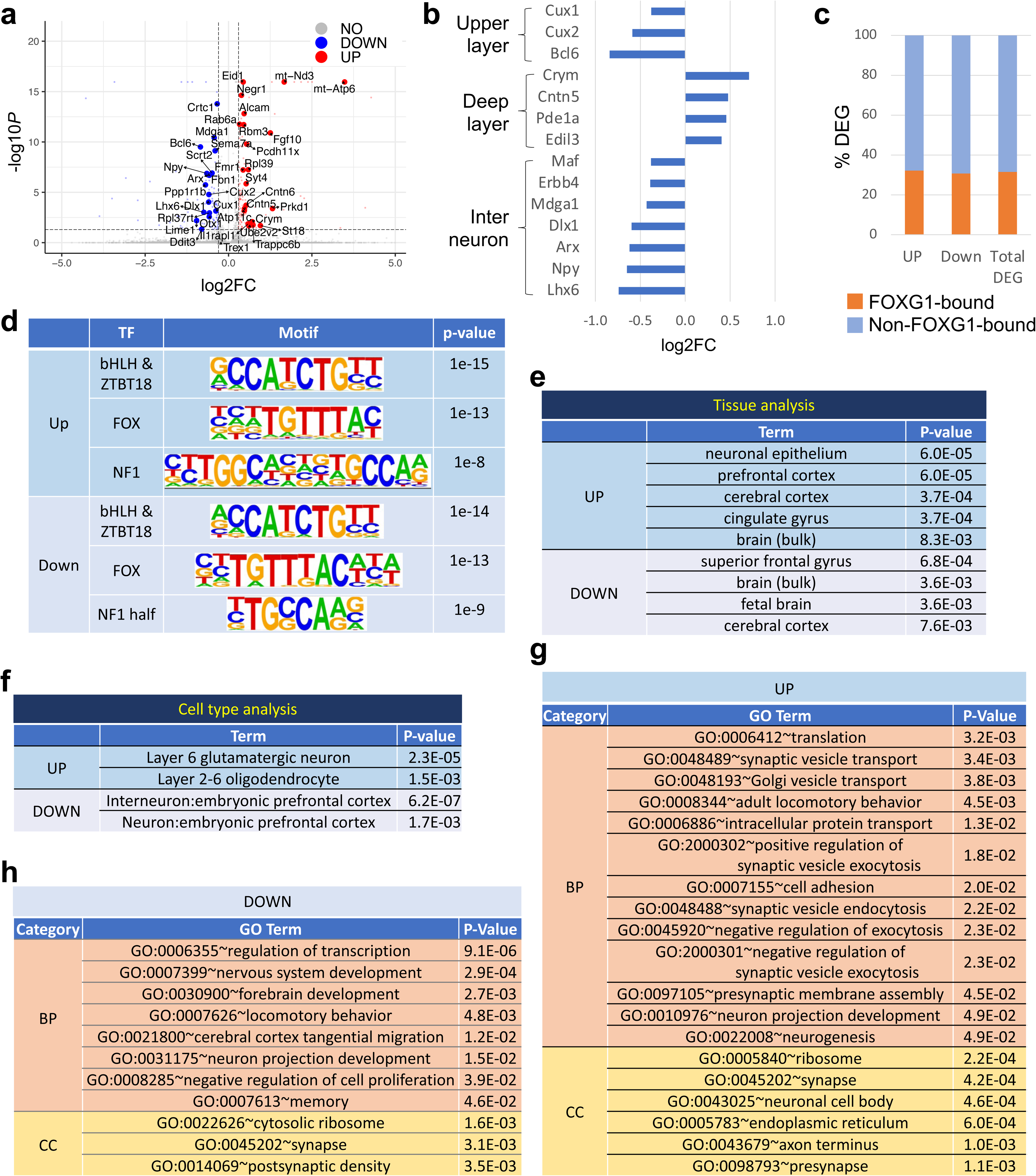
The dysregulated genes and pathways in Q84Pfs-Het cortex at P1. **a**, Differentially expressed genes (DEGs) in the RNA-seq analyses of P1 cortices of Q84Pfs-Het and WT mice as shown by the volcano plot (n=3 for Q84Pfs-Het mice; n=2 for WT). **b,** Cortical upper layer and interneuron genes were downregulated, whereas the deep layer genes were upregulated. **c,** The integration of DEGs of Q84Pfs-Het cortex and FOXG1 ChIP-seq data revealed the fraction of the DEGs that directly recruit FOXG1, as marked by orange. **d,** The motif analysis of FOXG1 ChIP-seq peaks associated with up- or down-regulated genes in Q84Pfs-Het cortex reveals the potential partner TFs that work with FOXG1 in cortex development. TF, transcription factor. **e-h,** Gene set enrichment analysis (GSEA) of DEGs. The tissue (**e**) and cell type (**f**) analyses show the tissue types and cell types that are associated with up- and down-regulated DEGs. Analyses of biological process (BP) and cellular component (CC) terms for up-regulated (**g**) and down-regulated (**h**) DEGs.

Approximately 30% of DEGs in Q84Pfs-Het cortex were associated with FOXG1-bound ChIP-seq peaks in the developing cortex^11^ regardless of up- or down-regulation (Fig. 5c), suggesting that FOXG1 regulates its direct target genes both negatively and positively. Consistently, the motif analysis^31^ revealed that the top two significantly enriched motifs in FOXG1-bound peaks annotated to either up- or down-regulated genes are the FOX motif and E-box. E-box serves as the binding site for basic helix-loop-helix (bHLH) and ZBTB18 (RP58) transcription factors, both of which were shown to collaborate with FOXG1^11,32^ (Fig. 5d). The NF1 site was also significantly associated with both up- and down-regulated FOXG1-target genes (Fig. 5d). These data suggest that a subset of dysregulated genes in Q84Pfs-Het mice are direct target genes of FOXG1 and that FOXG1 cooperates with ZBTB18, bHLH transcription factor, and NF1 in the developing cortex.

### The dysregulated pathways in Q84Pfs-Het cortex

To gain insights into molecular and cellular pathways leading to defects of Q84Pfs-Het mice, we performed the gene set enrichment analysis (GSEA) of DEGs using the Database for Annotation, Visualization, and Integrated Discovery (DAVID)^33^ and Enrichr ^34^.

In tissue analysis, DEGs were enriched in neuronal epitheliums, prefrontal cortex, cerebral cortex, cingulate gyrus, and superior frontal gyrus (Fig. 5e), consistent with the use of cortex for RNA-seq. In cell type analysis, the upregulated genes were significantly associated with cortical layer 6 glutamatergic neurons and, interestingly, oligodendrocytes, in which Foxg1 function remains unclear (Fig. 5f). The downregulated genes were most highly enriched for the “interneuron: embryonic prefrontal cortex” (Fig. 5f). Thus, our bioinformatic analyses suggest that the proportion of the three major cell types in the neonatal cortex, excitatory projection neurons, inhibitory GABAergic interneurons, and the oligodendrocyte lineage, was altered in Q84Pfs-Het mice.

Consistent with the known function of FOXG1 ^10–14^, DEGs of Q84Pfs-Het cortex were enriched for the genes controlling neuronal projection development, neuronal cell body, axon, and transcription regulations (Fig. 5g,h). Notably, the gene involved in the negative regulation of cell proliferation was downregulated in Q84Pfs-Het cortex (Fig. 5h), suggesting that NPCs may increase in Q84Pfs-Het cortex.

Intriguingly, many synaptic genes were significantly dysregulated in Q84Pfs-Het cortex. The upregulated genes were enriched for biological process (BP) terms of synaptic vesicle transport, synaptic vesicle exocytosis and endocytosis, and presynaptic membrane assembly, and cellular component (CC) terms of synapse and presynapse (Fig. 5g,h). Further, the downregulated genes were also strongly associated with the CC terms of synapse and postsynaptic density (Fig. 5g,h). Our results underline the new role of FOXG1 in synapse development. It is also noteworthy that the upregulated and downregulated genes were more strongly associated with presynapse and postsynaptic density, respectively.

Remarkably, both up- and down-regulated genes were enriched for translation and ribosome (Fig. 5g,h), suggesting defects in translation in Q84Pfs-Het brains. Other enriched terms, such as locomotory behavior and memory, are related to FS symptoms, including intellectual disability and hyperkinetic-dyskinetic movements ^19–23^.

Together, our genome-wide transcriptomics analysis indicates dysregulation in cell proliferation, axon projection, synapses, protein translation, and neuronal and glial cell compositions in Q84Pfs-Het cortex. Further, our results highlight the new role of FOXG1 in the oligodendrocyte development and synapse formation and function.

### Developmental defects of neural progenitors and excitatory and inhibitory neurons in Q84Pfs-Het cortex

We next assessed if the dysregulated genes in Q84Pfs-Het brains led to cortical defects suggested by our GO analyses, such as increased deep-layer glutamatergic neurons, reduced cortical interneurons, and downregulation of genes inhibiting cell proliferation and tangential migration of cortical interneurons (Fig. 5g-h).

Interestingly, although Q84Pfs-Het cortex was thinner than the WT cortex by E16, PAX6^+^ RGCs increased in Q84Pfs-Het cortex at E16 (Fig. 6a-c). Consistently, the number of phospho-histone H3^+^ proliferating cells in the VZ, especially at the ventricular surface, was significantly higher in Q84Pfs-Het cortex than in the control cortex (Fig. 6d,e). This increase aligns with the elevated levels of self-renewing RGCs with Q84Pfs expression in the developing cortex (Fig. 4) and stands in stark contrast to the reduced NPCs found in heterozygous mice carrying the *Foxg1*-null allele (*Foxg1^Cre/+^*, global Foxg1-Het) (Supplementary Fig. 3) ^2,5,8,9^. Together, our data demonstrate that RGCs increased in Q84Pfs-Het cortex.

**Fig. 6.**
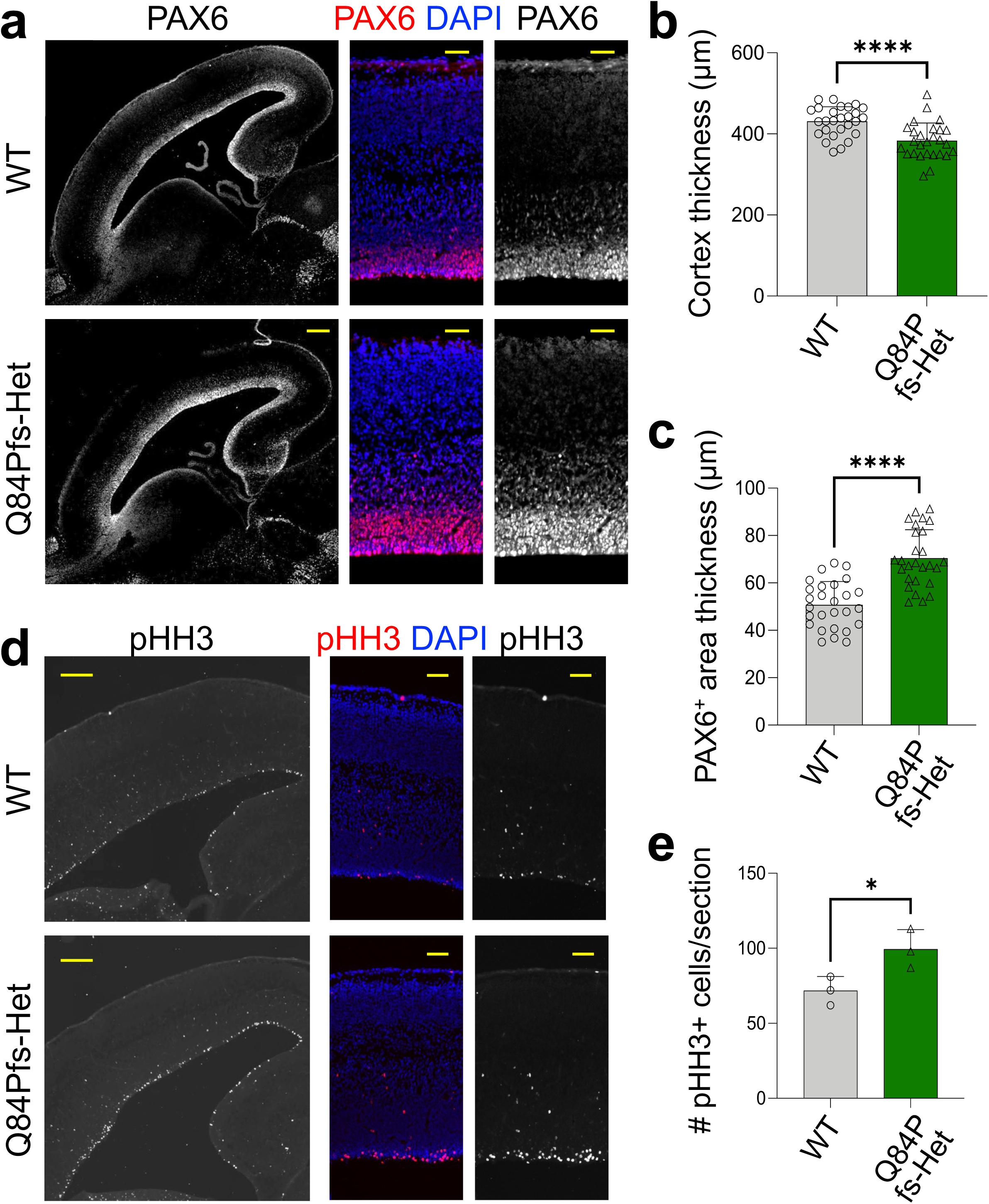
NPCs increased in Q84Pfs-Het cortex. The immunostaining analyses of Q84Pfs-Het and WT cortices at E16 with the NPC marker Pax6 (**a,c**) and the proliferation cell marker phosphorylated histone H3 (pHH3) (**d,e**). The quantification of cortex thickness (**b**), the thickness of Pax6^+^ progenitor areas (**c**), and the number of pHH3^+^ cells (**e**). Scale bars, 200μm (lower magnification images in **a, d**), or 50μm (higher magnification images in **a, d**). Thickness in (**b**,**c**) was measured in three independent areas per section (n = 3 mice/condition). The error bars (**b,c,e**) represent the standard deviation of the mean. \**p*<0.05 and \*\*\*\**p*<0.0001 in unpaired two-tailed t test. Only the representative images are shown.

The production of CUX1^+^ upper-layer neurons was delayed in E16 Q84Pfs-Het cortex, as shown by a lack of CUX1^+^ neurons in the superficial area of the cortex (Fig. 7a,b, Supplementary Fig. 4). CUX1^+^ upper-layer neurons remained significantly reduced in P1 and P30 Q84Pfs-Het cortex (Fig. 7c-h), indicating that the delayed upper-layer neuron generation is not compensated at the later stages. Notably, the number of BCL11B^+^ deep-layer neurons significantly increased, but the thickness of the deep-layer did not show a substantial change (Fig. 7a-i). These results suggest that during cortical neurogenesis, the production of deep-layer neurons increases at the expense of upper-layer neurons, and the reduction of the upper-layer neurons persists in the mature cortex in Q84Pfs-Het mice.

**Fig. 7.**
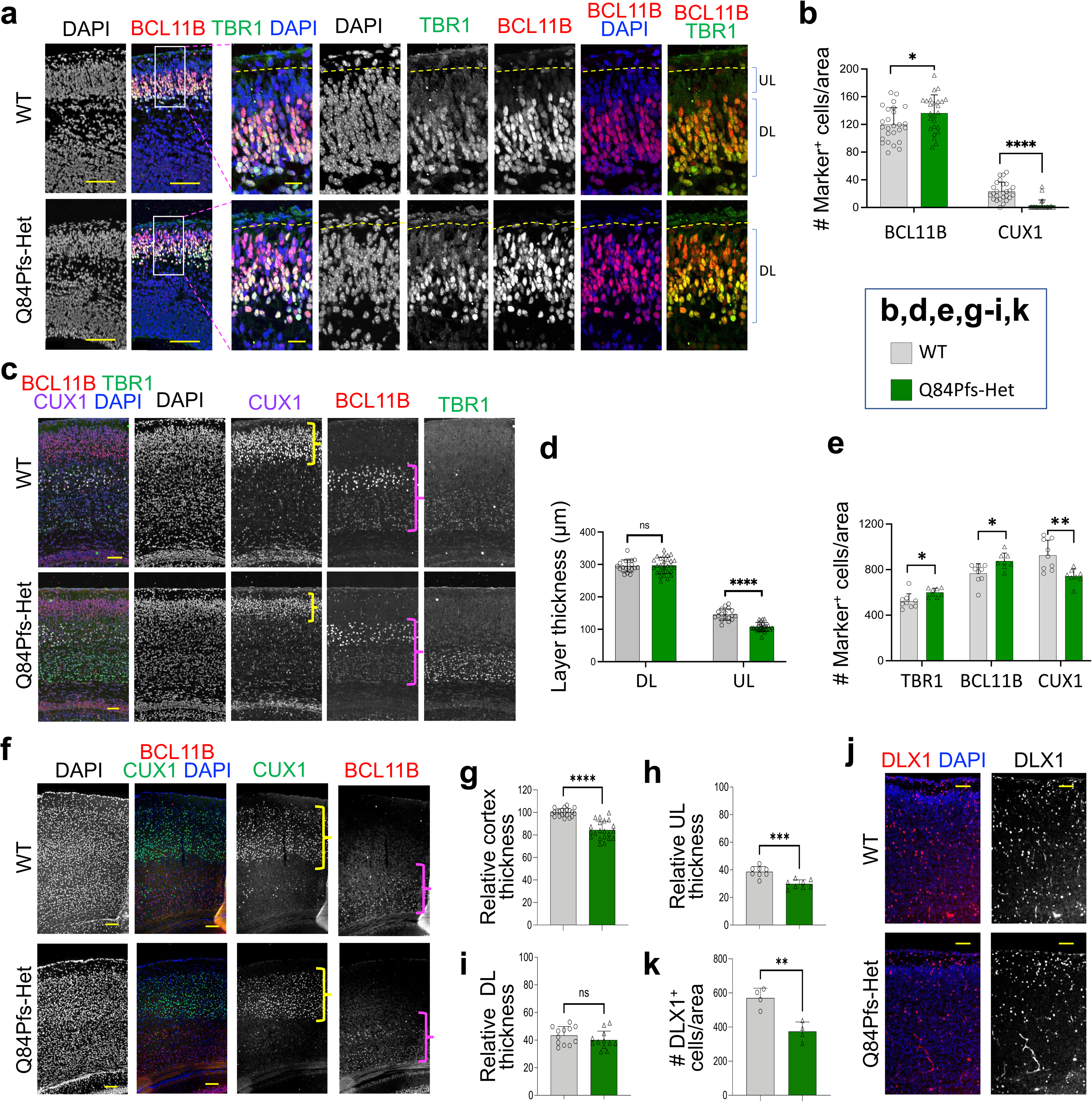
Q84Pfs-Het cortex showed a reduction of upper layer excitatory neurons and inhibitory interneurons and an increase of deep layer excitatory neurons. The immunostaining analyses of Q84Pfs-Het and WT brains with cortical excitatory and inhibitory neuronal markers at E16 (**a,b**), P1 (**c-e, j,k**), P30 (**f-i**). BCL11B and TBR1 mark deep layer (DL) neurons, whereas CUX1 labels upper layer (UL) neurons. DLX1 marks cortical interneurons. **a,b,** The yellow dotted lines indicate the upper limit of the cortical pyramidal neurons (**a**). BCL11B^+^ DL neurons increased, whereas UL neurons located above BCL11B^+^ neurons were markedly reduced in Q84Pfs-Het cortex at E16 (**a,b**). Similarly, the number of CUX1^+^ UL neurons, the thickness of CUX1^+^ UL, and the cortex thickness were significantly reduced in Q84Pfs-Het cortex at P1 (**c,e**) and P30 (**f,g,h**). The number of TBR1^+^ and BCL11B^+^ DL neurons increased, but the thickness of DL did not significantly change in Q84Pfs-Het cortex (**c-h**). The UL and DL were marked by yellow and magenta brackets, respectively (**c,f**). The number of DLX1^+^ interneurons was reduced in Q84Pfs-Het cortex (**j,k**). Scale bars, 100μm (lower magnification images in **a**), 20μm (higher magnification images in **a**), 500μm (**c,j**), or 1mm (**f**). n=4 mice/condition for **b,d,e,h,i,k**; n=7 mice/condition for **g**. The error bars represent the standard deviation of the mean. \**p*<0.05, \*\**p*<0.01, \*\*\**p*<0.001, \*\*\*\**p*<0.0001 and ns, non-significant in unpaired two-tailed t test. Only representative images are shown.

DLX1^+^ GABAergic interneurons were substantially decreased in Q84Pfs-Het cortex (Fig. 7j,k). While the overall level of cortical interneuron genes decreased in the RNA-seq dataset, their levels did not show significant changes when normalized by the expression of GAD1, a marker for GABA interneurons (Supplementary Fig. 5). Our data indicate that the number of inhibitory interneurons reduces in Q84Pfs-Het cortex.

### Axonal deficits in Q84Pfs-Het cortex

Q84Pfs-Het mice displayed two prominent axon deficits in the cortex, matching the dysregulation of neuron projection development genes in RNA-seq analyses (Fig. 5g,h). First, a substantial fraction of callosal axons was stalled at the midline and formed the Probst bundle in Q84Pfs-Het brains, leading to corpus callosum agenesis (Fig. 8a, Supplementary Fig. 6). Second, Q84Pfs-Het showed an increased number of aberrant L1^+^ axon bundles crossing the cortical plate toward the superficial area (Fig. 8b-d). These displaced L1^+^ axons expressed NTNG1 (netrin G1), a marker for axons derived from the thalamocortical neurons (Fig. 8b). As thalamocortical neurons do not express FOXG1, mislocated L1^+^NTNG1^+^ axons are likely to result from the changes in the cortex environment rather than cell-autonomous changes in the thalamocortical neurons.

**Fig. 8.**
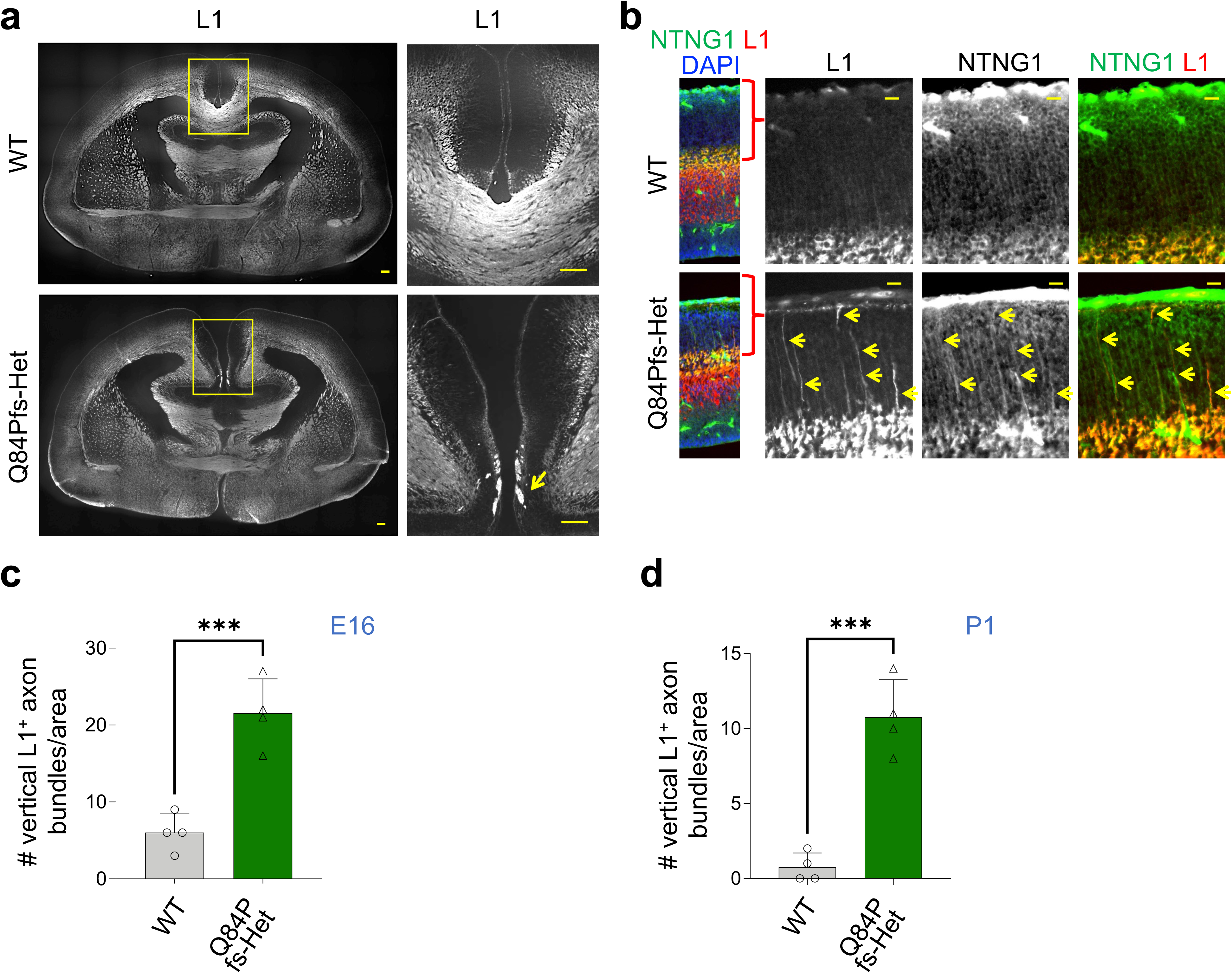
Q84Pfs-Het cortex exhibited axonal defects. The immunostaining analyses of Q84Pfs-Het and WT brains with the axonal marker L1 and thalamocortical axonal marker NTNG1 at P0 (**a**), E16 (**b,c**), and P1 (**d**). Probst bundles (yellow arrow in **a**) were formed at the midline of Q84Pfs-Het brains, but not of WT brains. In addition, Q84Pfs-Het cortex exhibited an increase in displaced L1/NTNG1-double positive axonal bundles (yellow arrows in **b**), as quantified (**c,d**). Scale bars, 1mm (**a**), or 200μm (**b**). n=4 mice/condition. The error bars represent the standard deviation of the mean. \*\*\**p*<0.001 in unpaired two-tailed t test. Only representative images are shown.

### Myelination defects in Q84Pfs-Het cortex

Although delayed myelination is one of the most common features of FS brains (Fig. 1d), the role of FOXG1 in the oligodendrocyte lineage development remains unclear. To test if the FS mouse model recapitulates this prominent white matter phenotype in humans, we monitored oligodendrocyte differentiation and myelination in Q84Pfs-Het mice at P30, the active myelination phase. Interestingly, OLIG2^+^ oligodendrocyte lineage cells, a majority of which represent OPCs, significantly increased in Q84Pfs-Het cortex (Fig. 9a,b), consistent with the association of upregulated genes with oligodendrocyte in the RNA-seq data (Fig. 5f). Despite the increased OLIG2^+^ cells, myelination was defective in Q84Pfs-Het brains, as shown by a significant decrease of MBP^+^ myelinated area and MBP immunostaining signals (Fig. 9c-e). Moreover, Q84Pfs-Het brains exhibited a marked reduction of the arborization and complexity of myelination patterns (Fig. 9c). Our results indicate that the oligodendrocyte development process was disrupted, and the myelination steps, including the microstructural organization of myelination, were impaired in Q84Pfs-Het brains, providing new insights into FS pathology.

**Fig. 9.**
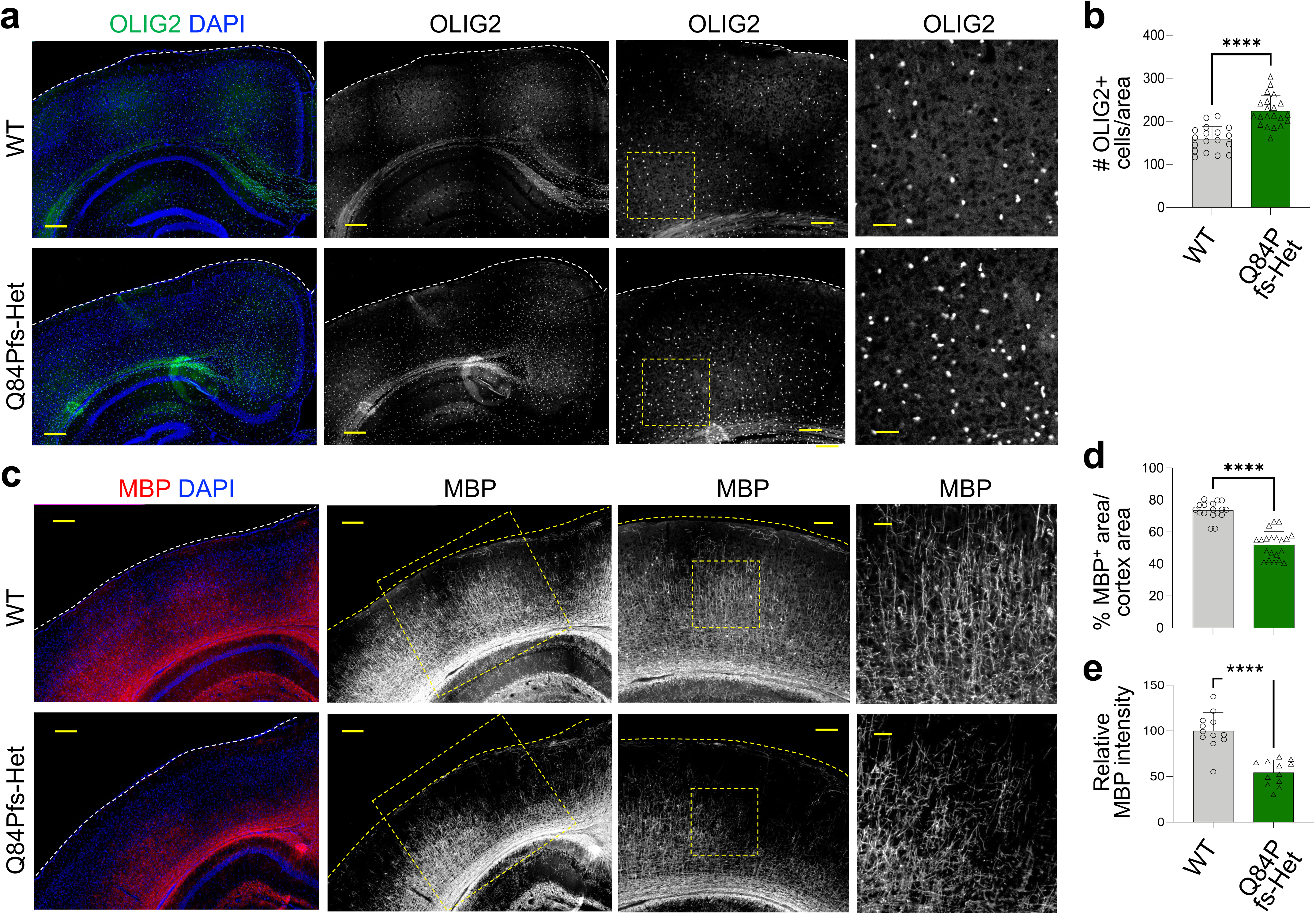
Q84Pfs-Het mice showed myelination deficits. The immunostaining analyses of Q84Pfs-Het and WT brains with oligodendrocyte lineage cell marker OLIG2 (**a,b**) and myelination marker MBP (**c-e**) at P30. Q84Pfs-Het cortex exhibited increased OLIG2^+^ oligodendrocyte lineage cells, markedly reduced myelinated areas, and disrupted patterns of myelination structure. The number of OLIG2^+^ cells was measured across three separate regions from the medial to the lateral cortex in coronal sections (**b**). The proportion of the myelinated area marked by MBP within the total cortical area (**d**) and the relative fluorescence intensity of MBP in the entire cortical region (**e**) were quantified. **b,d,e,** n=6 for WT; 7 mice for Q84Pfs-Het mice. The error bars represent the standard deviation of the mean. \*\*\*\**p*<0.0001 in unpaired two-tailed t test. Only representative images are shown. Scale bars, 1mm (the images in the first two columns in **a,c**), 300μm (the images in the third column in **c**), 500μm (the images in the third column in **c**), or 100μm (higher magnification images in **a,c**). The dotted rectangles marked the area that was magnified in the following columns.

### Dysregulated genes in Q84Pfs-Het cortex were associated with motor dysfunction and autistic-like behaviors

To ask if the dysregulated genes in the Q84Pfs-Het cortex are connected to specific human conditions, we performed GSEA with the DisGeNET database, which contains collections of genes and variants associated with human diseases ^35^ (Fig. 10a). The upregulated genes were significantly associated with impaired social interactions, Alzheimer’s disease, ASD, Parkinson’s disease, and autonomic nervous system disorders (Fig. 10a). The downregulated genes were enriched for dystonia, myoclonic encephalopathy, myoclonus, and ataxia, as well as major depressive disorder (Fig. 10a). Together, Q84Pfs-Het transcriptome changes are strongly linked to impairments in movement, social interactions, and autonomic nervous system function in Q84Pfs-Het mice.

**Fig. 10.**
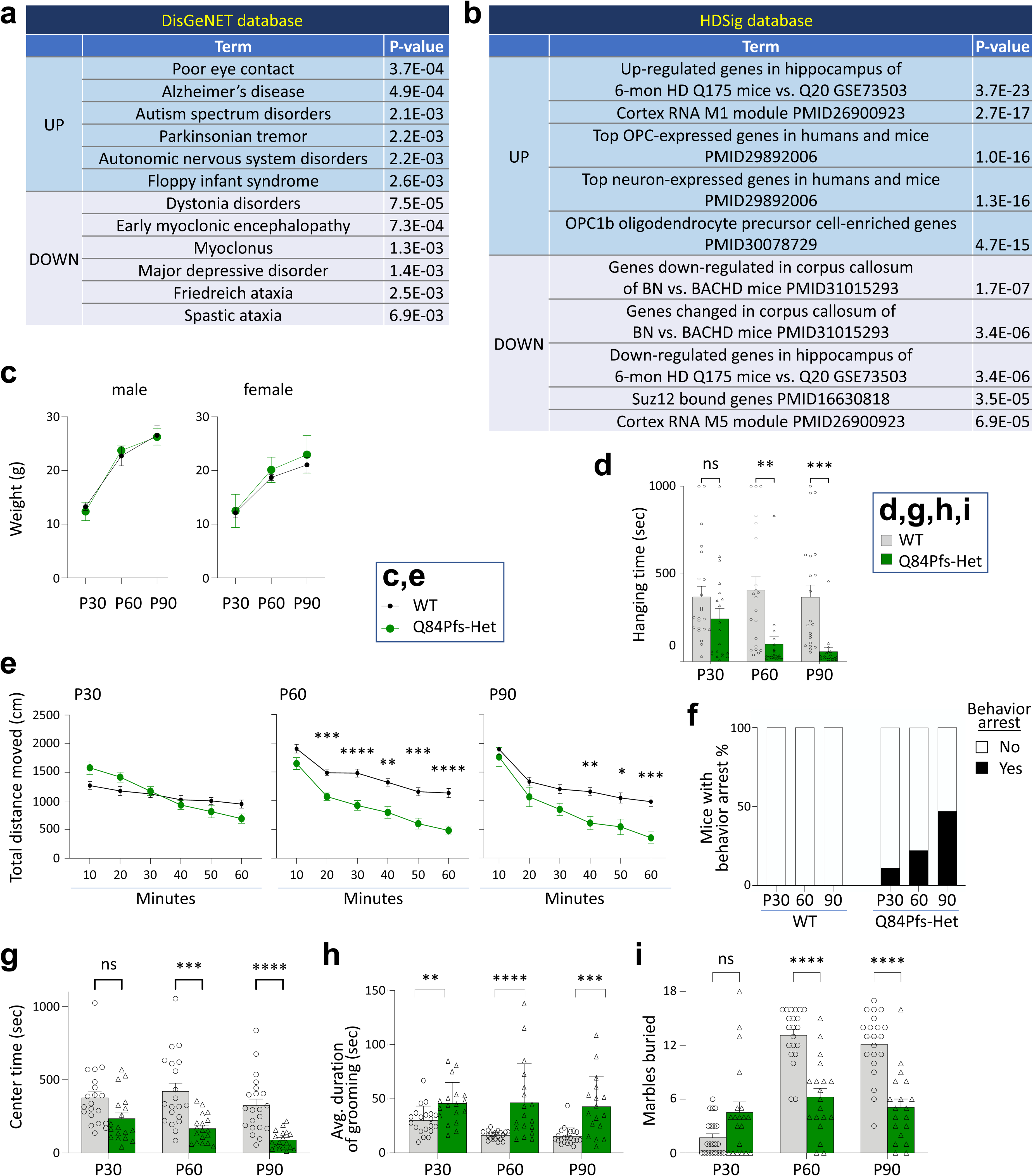
Q84Pfs-Het mice showed movement deficits, autism-like repetitive behaviors, and prolonged behavior arrest. **a,b,** Gene set enrichment analysis (GSEA) revealed the significant association of the DEGs in Q84Pfs-Het cortex with human disorders, such as autism (**a**), and Huntington’s disease mouse models and OPC genes (**b**). **c,** WT and Q84Pfs-Het mice showed no significant difference in body weights (n=WT 12 male, 9 female; Q84Pfs-Het 9 male, 10 female). **d,** Wire hanging test. Q84Pfs-Het mice showed a reduced hanging time. **e,** Open field test. Q84Pfs-Het mice showed reduced travel distance at P60 and P90, but not at P30. **f,** Some Q84Pfs-Het mice displayed extended periods of behavioral arrest, characterized by paused movement lasting more than 3 min at one episode during a one-hour-long open field test. In contrast, none of the WT mice showed such prolonged behavior arrest. The black bars represent the percentage of mice that experienced behavioral arrest out of the total number of mice assessed. **g,** Center time analysis during a one-hour-long open field test. Q84Pfs-Het mice showed a reduced center time at P60 and P90. **h,** Q84Pfs-Het mice exhibited significantly increased self-grooming behavior relative to WT, as measured by the average duration of an individual grooming event. **i,** Q84Pfs-Het mice buried significantly fewer marbles than WT mice at P60 and P90 but showed a tendency to bury more marbles at P30. **c-i,** Error bars, SEM. \**p*< 0.05, \*\**p*<0.01, \*\*\**p*<0.001, \*\*\*\**p*<0.0001 and ns, not significant in the two-way ANOVA test (**c,d,e,g,i**) or the Mann-Whitney Test (**h**). Each circle and triangle correspond to one animal. **c,** WT n = 22 (12 male, 10 female), Q84Pfs-Het n= 20 (10 male, 10 female). **d,i,** For P30, WT n = 22 (12 male, 10 female), Q84Pfs-Het n= 20 (10 male, 10 female); for P60 and P90, WT n = 21 (12 male, 9 female), Q84Pfs-Het n = 19 (9 male, 10 female). **e-h,** For P30 and P60, WT n = 20 (12 male, 8 female), Q84Pfs-Het n = 18 (8 male, 10 female); for P90 in **e,g**, WT n = 21 (12 male, 9 female), Q84Pfs-Het n =18 (8 male, 10 female) and for P90 in **f,h**, WT n = 21 (12 male, 9 female), Q84Pfs-Het n = 17 (7 male, 10 female).

Given that both FS and Huntington’s disease (HD) exhibit many neurological symptoms, including abnormal involuntary movements ^22,36^, we compared the dysregulated genes between Q84Pfs mice and HD using the database of HD molecular signatures (HDSigDB, hdinhd.org) (Fig. 10b). This analysis revealed the striking resemblance between Q84Pfs-Het mice and HD mouse models ^37,38^. The high-ranking categories that resemble the up- and down-regulated genes in Q84Pfs-Het mice were the up- and down-regulated genes in the hippocampus of the HD mouse model Q175 ^37^, respectively (Fig. 10b). Notably, the two categories among the top five gene sets for the upregulated genes were oligodendrocyte progenitor cell (OPC) genes (Fig. 10b), consistent with our finding that Q84Pfs-Het cortex showed the increased OLIG2^+^ oligodendrocyte lineage cells (Fig. 9a,b).

Collectively, the transcriptome changes in Q84Pfs-Het cortex are closely connected to the clinical features of FS, such as movement disorders, autism-like behaviors, and social impairment.

### Q84Pfs-Het mice showed movement deficits, repetitive behaviors, and prolonged behavior arrest

To investigate whether Q84Pfs-Het mice exhibit behavior phenotypes corresponding to the above molecular and cellular changes, we performed behavioral assessment tests at adolescence stage P30 and adult stages P60 and P90.

There was no significant difference in body weights of Q84Pfs-Het and WT mice at these stages (Fig. 10c). In the wire hang test that evaluates the motor function and muscle strength, Q84Pfs-Het mice showed significantly reduced hang time, which worsened as mice aged (Fig. 10d). In the open field test that monitors locomotor activity and exploratory behaviors, Q84Pfs-Het mice moved significantly less than WT mice at P60 and P90 (Fig. 10e). At P30, Q84Pfs-Het mice showed a tendency for travel distance reduction over time compared to WT mice, but the total travel distance did not differ significantly (Fig. 10e). Q84Pfs-Het mice exhibited a strikingly extended behavior arrest, defined by paused locomotion for longer than 3 min at one episode during free exploration of the arena (Fig. 10f, Supplementary Fig. 7 and videos). Throughout these episodes of remarkably pronounced stops, Q84Pfs-Het mice showed a high degree of postural control, typically positioning their bodies facing the center of the arena, and no apparent overt movements, such as visual survey, rearing, or digging. The proportion of Q84Pfs-Het mice that exhibited the extended behavior arrest was 11%, 22%, and 47% at P30, P60, and P90, respectively, whereas WT mice did not show behavior arrest without purposeful movements under the same condition (Fig. 10f). Notably, Q84Pfs-Het mice that do not present prolonged behavior arrests also showed a decreased travel distance. Thus, movement reduction cannot be attributed solely to the extended arrest.

Next, we assessed ASD-related behaviors. The center time in the open field test was markedly decreased as Q84Pfs-Het mice aged from P30 to P90, relative to WT mice (Fig. 10g), indicating increased anxiety levels. Q84Pfs-Het mice exhibited a significant increase in repetitive grooming behavior at all three stages, with a tendency for more significant differences from WT mice as they age (Fig. 10h). These results suggest heightened repetitive behavior in Q84Pfs-Het mice. Intriguingly, in the marble-burying test, Q84Pfs-Het mice buried more marbles at P30 but fewer marbles at the adult stages than WT mice at P60 and P90 (Fig. 10i), suggesting age-dependent changes in the marble burying behavior.

Together, our data uncovered that Q84Pfs-Het mice present FS-like motor deficits, such as poor motor coordination. In addition, Q84Pfs-Het mice display ASD-related behaviors, such as increased anxiety levels, and repetitive behaviors. All of these behavioral traits have been observed in FS ^19–23^ and are strongly linked to molecular and cellular changes in Q84Pfs-Het brains.

## Discussion

Most FS patients possess mutations within the *FOXG1* gene coding region, which are likely to produce faulty FOXG1 protein products that impact disease mechanisms and progression. Therefore, the currently available *Foxg1*-null mouse lines are inadequate for investigating FS pathophysiology because they delete the entire FOXG1-coding region ^2,9,26,39,40^. Here, we established Q84Pfs-Het mice as the first patient-specific FS mouse model that accurately replicates human genetic conditions of FS. Remarkably, the heterozygous mice carrying only a single allele of *Q84Pfs* variant, without creating the homozygous conditions, recapitulated a wide range of human FS symptoms, such as brain structural deficits and ASD-like behaviors. Using this new FS mouse model, we uncovered molecular and cellular changes, such as oligodendrocyte lineage deficiency and dysregulation of synaptic genes, leading to the key hallmarks of FS, including delayed myelination, movement deficits, and ASD-like behaviors. Furthermore, we found that Q84Pfs, the N-terminal fragment of the FOXG1 protein, is expressed in Q84Pfs-Het brains. Intriguingly, the Q86Pfs fragment (the human counterpart of the mouse Q84Pfs) was found to associate with FOXG1 full-length protein and inhibit neuronal migration while promoting a self-renewing RGC fate. This suggests that the defective FOXG1 protein product plays a significant role in the haploinsufficiency disorder FS.

In contrast to many ASD mouse models that typically involve the complete knockout of ASD-related genes, which does not accurately reflect the heterozygous or epigenetically modified nature of these genes in human conditions, our study with Q84Pfs-Het mice closely mirrors human FS at a genetic level, offering substantial benefits for understanding and treating neurodevelopmental disorders. The Q84Pfs-Het model led to the identification of dysregulated genes in the brains of mice with FS-like conditions, which likely contribute to cellular, structural, and behavioral abnormalities. Furthermore, the model has revealed novel mouse phenotypes that closely align with the clinical manifestations observed in human FS, such as impairments in oligodendrocyte maturation and myelination, heightened repetitive behaviors, and anxiety. Therefore, the Q84Pfs-Het mouse serves as an exemplary model to decipher the pathogenic mechanisms underlying these phenotypes and their equivalent human symptoms. Finally, the *Foxg1* gene heterozygous condition of Q84Pfs-Het mice presents a unique opportunity to test therapeutic strategies that would not be viable in homozygous models. This includes strategies aimed at enhancing FOXG1 expression from the WT allele, thereby increasing the levels of functional FOXG1 protein.

Our concurrent molecular and behavioral examinations of Q84Pfs-Het mice have yielded critical insights into the development of FS. The level of *Foxg1* mRNA in Q84Pfs-Het mice did not diminish (Supplementary dataset), indicating that the mRNA from the *Q84Pfs-Het* allele was not subject to nonsense-mediated decay. Indeed, Q84Pfs-Het brains expressed the Q84Pfs protein (Q86Pfs protein in humans), the N-terminal fragment of the FOXG1 protein with unique amino acid signatures of histidine, proline, and glutamine repeats. Notably, Q84Pfs forms the pronounced speckles in HEK293 cells and cortical progenitors and neurons, features distinct from FOXG1 full-length protein. Further, Q84Pfs associates with FOXG1 full-length protein and recruits FOXG1 into speckle domains. These molecular characteristics of Q84Pfs suggest that Q84Pfs can interfere with action of wild-type FOXG1 via mechanisms such as sequestering FOXG1 from its genomic target loci. In the developing cortex, Q86Pfs protein drives the RGC fate with enhanced self-expanding capabilities and suppresses neuronal migration. Thus, the presence of Q84Pfs fragment is likely to contribute to the phenotypic differences between Q84Pfs-Het and the existing global *Foxg1*-Het mice^41^ in the developing cortex. Q84Pfs-Het cortex showed the gene expression changes that predict various movement deficits and ASD-like behaviors. Consistently, adult Q84Pfs-Het mice exhibited the corresponding behavioral characteristics. This close correlation between gene expression anomalies and behavioral traits highlights specific genes whose dysregulation may be responsible for movement and ASD-like symptoms in Q84Pfs-Het mice. *Fmr1*, *Nalcn*, *Il1rapl1*, and *Slc16a2*, which are aberrantly upregulated in Q84Pfs-Het brains, may be involved in intellectual disability, social impairment, and autism-like behaviors, given that these genes are associated with neurodevelopmental disorders with the described signature behaviors ^42–45^. A subset of FS symptoms, such as dystonia, myoclonus, and ataxia, may be attributed to the changes of *PLEKHG2*, *DST*, *ARX*, *AAAS*, *KCTD17*, *PRDM8*, and *CRTC1* in FS brains as these genes are downregulated in Q84Pfs-Het brains and play a role in these movement and muscle coordination ^46–52^. The remarkable similarity of dysregulated genes between Q84Pfs-Het and HD mouse models suggests that the shared gene expression changes are responsible for common symptoms between FS and HD, such as involuntary jerky and writhing movement. Our studies also revealed that the genes involved in synaptic vesicle exocytosis, endocytosis, and synapse organization and membrane assembly are dysregulated in Q84Pfs-Het cortex, strongly implying synapse deficiency as a mechanism driving FS symptoms, including prolonged behavior arrests and cognition impairments. Of note, the prolonged behavior arrests similar to the behavior of Q84Pfs-Het mice have been described for seizures, like focal behavior arrest seizures (aka, focal impaired awareness seizures, hypokinetic seizure, localized seizure with behavior arrest, or complex partial seizures) ^53–56^. As this type of seizure has been observed in FS ^57^, it will be interesting to perform electroencephalogram (EEG) recording paired with behavioral assessment in Q84Pfs-Het mice and examine if behavior arrest is correlated with specific types of EEG activities. Given the symptomatic similarity between FS and other neurological disorders, the pathogenic mechanism studies in Q84Pfs-Het mice provide significant insights into a broader range of neurodevelopmental disorders, including ASD.

FOXG1 plays cell type-specific and developmental timing-dependent roles in forebrain development, at least via the two mechanisms. First, the dynamic expression of FOXG1 plays a key role. The downregulation of FOXG1 during neurogenesis is crucial for the cell cycle exit and differentiation of neural progenitors ^8,9,58,59^. After neuronal differentiation, FOXG1 is upregulated in postmitotic neurons, in which FOXG1 promotes neuronal morphological transition and migration ^10,11^. Second, FOXG1 partners with other transcription factors in a cell context-specific manner in selecting target genes. Our motif analysis of FOXG1-bound genomic loci of DEGs identified E-box, the binding sites for FOXG1’s partner transcription factors ZBTB18 and NEUROD1 ^11,32^, along with the FOX motif. Our data integrating RNA-seq and ChIP-seq datasets also suggest that NF1 collaborates with FOXG1 in the cortex. Notably, NF1 is a family of closely related transcription factors, NF1A, NF1B, NF1C and NF1X, and in particular, NFIA and NFIB have been found to be involved in haploinsufficiency syndromes that share some features with the FS, such as corpus callosum agenesis and intellectual disability^60–62^.

It is noteworthy that our western blot analysis of HEK293 cells expressing FOXG1 full-length did not detect fragments of the FOXG1 protein, contrasting the report that the cleaved product of FOXG1 localizes in mitochondria ^63^. Given the use of the different cells between the two studies, specific cell context may be necessary for FOXG1 protein cleavage, which is an interesting topic for future studies.

The myelination deficits in Q84Pfs-Het mice suggest a key role of FOXG1 in oligodendrocyte development. In Q84Pfs-Het brains, OLIG2^+^ oligodendrocyte lineage cells increased but myelination was impaired, suggesting FOXG1 functions at multiple points of oligodendrocyte development. Combined with the finding that overexpression of FOXG1 inhibits the generation of OPC from neural progenitors^64^, our results raise the possibility that *FOXG1* haploinsufficiency triggers the overproduction of OPCs from neural progenitors but inhibits differentiation and maturation of oligodendrocytes, thereby resulting in accumulated OPCs and reduced myelination. It is also possible that the rise in OPCs may be a compensatory response to myelination deficits ^65^. Given that proper nerve myelination is crucial for movement and various brain functions, and considering that myelination mainly occurs postnatally and continues into young adulthood in humans ^66^, targeting dysmyelination represents a viable therapeutic approach for FS as exemplified by the AAV-FOXG1 approach ^67^.

Collectively, our study established Q84Pfs-Het mice as the FS mouse model that will expedite our understanding of the pathophysiology of FS and serve as an essential platform for therapeutic development. Importantly, our study uncovered new molecular and cellular mechanisms of FOXG1 function and FS etiology.

## Methods

### FOXG1 Syndrome Research Registry & Ciitizen Databank

The International FOXG1 Syndrome Research Registry is an online platform that collects information from caregivers of children and adults with FS, including results of genetic testing, clinical phenotypes, and developmental outcomes [https://foxg1.beneufit.com/login.php]. Ciitizen®, a wholly owned subsidiary of Invitae Corporation, is a patient-centric real world data platform that transforms medical records into structured datasets that describe clinical phenotypes and medical interventions [https://www.medrxiv.org/content/10.1101/2023.03.02.23286645v1]. In partnership with the FOXG1 Research Foundation, >120 individuals with a FOXG1 variant have enrolled in Ciitizen. The sex of human participants was annotated from the participant’s birth certificate. As FOXG1 syndrome is a rare disorder, recruitment for the FOXG1 syndrome patient registry and natural history study is broad and does not select one sex/gender. In brief, the platform supports medical records that are collected on behalf of participants through the Health Insurance Portability and Accountability Act (HIPAA). Following record collection, each medical document is systematically reviewed for relevant data variables, including presenting diagnoses and diagnostic imaging results. Genetic variants were extracted from clinical genetic test reports; loss-of-function variants in FS, including frameshift variants, are known to be pathogenic. To support harmonization of disparate data sources, extracted phenotypic data were mapped to SNOMED CT, an internationally recognized terminology (US Edition, version 2022_03_01) [www.snomed.org]. The resulting data elements were stored securely in HIPAA-compliant, controlled access, indexed database.

Caregivers and/or legal guardians of study participants provided broad consent to share de-identified data for research. The generation and subsequent analysis of participant data used in this study received determinations of exemption through a central institutional review board, Pearl IRB. Reasonable requests for data access can be made to.

### Animals

All experimental procedures were approved by the IACUC at University at Buffalo and performed in accordance with the NIH Guide for the Care and Use of Laboratory Animals. Q84Pfs mutant mice were generated with the CRISPR/Cas9 gene editing system using guide RNA (CCCAGCAGCCGACGACGACA) and single stranded oligonucleotides (ssODN) containing the mutation c.250dupC to introduce the Q84Pfs allele and the restriction enzyme site (HindIII) for genotyping (T CCA CCG CCG CCC CAG CAG CAG CAG CAG CAG CCG CCC CCG GCC CCG CAG CCC CCG C CAG GCG CGC GGC GCC CCA GCA GCC GAC GAC GAC AAG CTT CCC CAG CCG CTC CTG CTC CCG CCC TCC ACC). Potential founder mice were genotyped by Sanger sequencing and restriction enzyme digestion for PCR product surrounding the target site. The targeted founder mice from CRISPR/Cas9 microinjection were backcrossed to C57BL/6J mice for at least five generations to eliminate potential off-target mutations before any experiments.

### Behavioral tests

Mice were maintained on a 12 h light/dark cycle, and all behavioral tests were performed during the light phase. All animals were allowed to habituate in the behavior testing room (with attenuated light and sound) for at least 30 min before the tests. All arenas and objects used in behavioral analysis were cleaned with Alconox and then distilled water between test sessions.

*<u>Open field test</u>*: The open field test was utilized to examine locomotor activity and to monitor anxiety-like behaviors as well as repetitive behaviors such as jumping and self-grooming. Mice were placed into a Plexiglas chamber with transparent walls and white floor (40cm W X 40cm D X 30cm H) and then allowed to move freely for 60min. All activity was monitored by SuperFlex Open Field system (Omnitech Electronics) and the total moved distance and the time spent in the center (20 cm X 20 cm within the center of the floor) were collected and analyzed by Fusion software (AccuScan Instruments Incorporated). *<u>Grooming</u>*: Grooming was assessed manually using video recordings from the open field test. Each mouse was monitored for 60 minutes. For a grooming event to count, the mouse had to groom for at least 2 seconds. If the mouse stopped grooming, but resumed in 5 seconds or less, it was counted as the same grooming event. If the mouse moved to a new location in the cage and continued grooming, even if it resumed in 5 seconds or less, it was counted as a new grooming event. *<u>Prolonged behavior arrest</u>*: During the open field test, prolonged behavior arrest was marked whenever a mouse remained sitting in one location in the cage, largely unmoving, for at least 3 minutes. During this time, the mice did not appear to perform any purposeful movement, such as grooming, rearing, digging, or other exploratory behavior. *<u>Wire hanging test</u>*: Mice were placed inside a basket with wire grids on the bottom, and the basket was slowly inverted to allow the animal to hang from the grid. All sessions were video-recorded, and the hanging time was analyzed manually. The hanging time was defined from the time the basket was inverted to the time that the mouse fell off the wire grid. Mice were tested on three consecutive trials with at least 5 min intervals between trials. All trials ended upon reaching 1000 seconds. *<u>Marble burying</u>*: Mice were gently placed in the periphery of testing cages (arena size: 30 cm × 20 cm × 13 cm, corncob bedding depth: 5 cm) containing 18 glass marbles (evenly spaced with three rows of six marbles on the top of bedding), and then the cages were covered with the transparent top (height 13 cm). Mice were carefully removed from the cages, and the number of marbles buried was recorded at the end of the 30 min exploration period. Marbles were counted as buried when they have covered at least 2/3 depth with bedding.

### Brain preparation, immunohistochemical analysis, and image acquisition

Dissected brains were fixed in 4% paraformaldehyde in phosphate-buffered saline (PBS) at 4°C overnight, equilibrated in 30% sucrose, and embedded in Tissue Freezing Medium^TM^ (Electron Microscopy Sciences). Frozen sections were cut with 8µm (Fig. 3a, Fig. 7a,c, and Supplementary Fig. 3), 15µm (Fig. 3f, Fig. 6, Fig. 8, and Supplementary Fig. 4) or 18µm (Fig. 7f,j, and Fig. 9) using cryostat (CM1950, Leica). Position matched sections were then stained following the standard immunohistochemical method. Sections were incubated with the primary antibody and the secondary antibodies conjugated to fluorophores (Jackson ImmunoResearch), and then counterstained with DAPI to reveal nuclei. Images were collected through DM6 fluorescence microscope (analyzed by Leica Software LasX) or Eclipse Ti2 confocal microscope (analyzed by Nikon Software). The primary antibodies included rabbit anti-OLIG2 (Millipore AB15328, 1:1000), rat anti-L1 (NCAM) (Millipore MAB5272, 1:1000), rat anti-CTIP2 (Abcam AB18465, 1:1000), rat anti-MBP (Millipore MAB386, 1:500), rabbit anti-CUX1 (Santacruz sc-13024, 1:500), chicken anti-TBR1 (Millipore AB2261, 1:1000), rabbit anti-PAX6 (Biolegend Poly19013, 1:1000), rabbit anti-phospho-Histone H3 (Ser10) (Cell Signaling #9701S, 1:1000), rabbit anti-DLX1 (Chemicon AB5724, 1:1000), goat anti-NETRIN-G1a (R&D AF1166, 1:500), and rat anti-BrdU (Abcam AB6326, 1:1000) antibodies. The secondary antibodies from Jackson ImmunoResearch Laboratories included donkey anti-rat Cy3 (Cat.712-165-153, 1:500), donkey anti-rabbit Alexa Fluor 647 (Cat.711-605-152, 1:500), donkey anti-rabbit Alexa Fluor 488 (Cat.711-545-152, 1:500), donkey anti-chicken Alexa Fluor 488 (Cat.103-545-155, 1:500) and goat anti-rat Alexa Fluor 594 (Cat.112-585-167, 1:500) antibodies.

Image data were collected through DM6 fluorescence microscope and analyzed by Leica Software LasX. To measure OLIG2^+^ cells, 3 areas were selected in each section of the cortex. One Region of Interest (ROI) (x = 1 mm; y = cortex thickness) per each area immunostained with anti-OLIG2 was counted for the number of positive cells.

High-resolution confocal images (60X magnification, 2 X 4 tiles) were employed in CellProfiler ^68^ for the automated quantification of immunofluorescent intensity in each cortex cell. The following pipeline was implemented: The Crop module was utilized to eliminate unnecessary areas in the images. The Identify Primary Objects module, employing Global thresholding, the Otsu approach, and based on object intensity settings, was employed to recognize the FOXG1 antibody immunostaining signal in each individual cell. The settings were adjusted to capture all positive signaling while minimizing noise. The Measure Object Intensity module was used to generate intensity data for further analysis. The Export to Spreadsheet module was applied to export all data into Excel format for subsequent processing.

### Nissl staining

Slides with cryosectioned frozen sections (18µm) were incubated in ethanol/chloroform mixture, mixed 1:1 volumetrically, at RT overnight. Incubated slides went through rehydration steps in three steps, 100% ethanol, 95% ethanol and then distilled water for 10 minutes each. After rehydration, slides were stained in warm 0.2% cresyl violet solution for 12 min, followed by brief washing and then incubating in 95% ethanol, 100% ethanol and xylene for 20 minutes each for differentiation, dehydration, and clearing, respectively. Coverslip was mounted with Permount^TM^ Mounting Medium (Fisher Scientific). Images from stained specimen were acquired by Axio Scan.Z1 (Zeiss) under bright field with a 10X objective.

### *In utero* electroporation

Male and female C57BL/6J mice purchased from Jax (Strain #: 000664) were used for breeding and timed-pregnant females (E15.5) were anesthetized with 2% isoflurane infused into 2L/min oxygen. The females were placed on the heat pad to prevent hypothermia during procedure. Embryos were exposed after making 1.5 cm incision through midline of pregnant female’s abdomen. pCIG2-Ires-EGFP or pCIG2-3XFLAG-Q86Pfs-Ires-EGFP (1.5 µg/ µL) was injected into the lateral ventricle and electroporated with Tweezertrodes across the cortex using electroporation system (BTX Harvard Apparatus, ECM830, 45-0052). For electroporation, 5 trains of pulses of 50 ms long 45V shock were given with interval of 950 ms between pulses. Embryos were then put back into dam’s abdominal cavity and incision was sutured with non-absorbable Nylon thread. Brains were harvested from embryos two or three days after electroporation. To label cell cycle S phase cells, BrdU (50mg / kg body weight) was administered by intraperitoneal injection 2 hours before brain harvest.

### RNA-seq data analyses

RNA-seq analyses were performed using P1 cortices of Q84Pfs-Het and WT mice (n=3 for Q84Pfs-Het mice; n=2 for WT). The appropriate sample size was validated using ssizeRNA, which calculates the sample size while controlling FDR for RNA-seq experimental design^69^, with the default values except for the fold change and the proportion of non-differentially expressed genes (nonDEGs). This analysis showed that the sample size five has discriminative power. FastQC was used to check the quality of the raw RNA-seq data. Clean reads were aligned to the mouse reference genome (mm10) using STAR (v2.6.1d) ^70^ with default parameters and the gene expression levels were quantified using RSEM (v1.3.3) ^71^. The DEGs were identified using EBseq ^72^ based on the expected counts. The absolute value of log2 fold change of 0.3 and false discovery rate (FDR) of 0.05 were set as criteria to determine if the expression level of a gene was significantly changed. In this manner, 222 DEGs including 118 upregulated and 104 downregulated DEGs were identified in total. We subsequently carried out GSEA using DAVID and Enrichr on the upregulated DEGs and the downregulated DEGs, separately.

### Transfection, coIP assays, and immunohistochemical analysis in HEK293 cells

The FOXG1-fl- and Q86Pfs-coding genes were subcloned into the pCIG vector containing IRES-GFP sequences. The mammalian expression vectors encoding the FOXG1-fl and Q86Pfs, with or without epitope tags, were prepared using Qiaprep spin miniprep kits (Qiagen, 27106). DNA transfections into HEK293T cells were performed using TurboFect Transfection Reagent (Thermo Scientific R0531) following the provided manuals. For coIP studies testing the association between FOXG1-fl proteins, 10 cm dishes of HEK293 cells were transiently transfected with DNA constructs coding for Flag-FOXG1-fl (2.5µg) and HA-FOXG1-fl (2.5µg). To investigate the interaction between FOXG1-fl and Q86Pfs, 10 mm dishes of HEK293 cells were transiently transfected with DNA constructs coding for Flag-FOXG-fl (2.5µg) and Q86Pfs (2.5µg, no epitope tag). For cellular immunostaining, 8-well chamber slides (BD Falcon, 354118) of HEK293T cells were transiently transfected with DNA constructs coding for FOXG1-fl (375ng) + empty vector (375ng), or Q86Pfs (375ng) + empty vector (375ng), or FOXG1-fl (375ng) + Q86Pfs (375ng). The cells, with the transfection reagent/DNA mixture, were incubated at 37 °C in a CO_2_ incubator for 24 hours before coIP or immunostaining.

For coIP assays, lysis of 10 cm HEK293 cells was carried out using 500μl PBS supplemented with 5mM EDTA, 0.02% sodium azide, and a protease inhibitor cocktail (IP buffer). Following sonication for 30 seconds, the cells were incubated in IP buffer with rotation at 4 °C for 1 hour. Subsequently, the lysate was centrifuged at 13,000 rpm for 10 minutes at 4 °C, and 20μl of the supernatant was saved as input. The remaining supernatant was transferred to a new tube, and 20μl beads were added for a 1-hour pre-clear incubation. The supernatant was then divided into two tubes, and IP buffer was added to achieve a final volume of 500μl. To each tube, 20μl of protein A/G-conjugated beads was added, followed by an overnight incubation at 4°C with 2μl of anti-Flag antibody (Sigma, F1804) or IgG (cell signaling, #3900). After discarding the supernatants, the beads were washed 3-5 times with IP buffer. Finally, 2X Laemmli sample buffer (Bio-RAD, #1610737) was added, and the mixture was boiled for 8 minutes before conducting western blot analysis using anti-HA antibody (Sigma H6908) or anti-FOXG1 antibody (Invitrogen #PA5-41493).

For immunostaining analysis, cells were washed twice in PBS and fixed in 4% PFA for 15 minutes. Subsequently, they underwent three PBT washes to eliminate residual PFA. The cells were then incubated in 0.5% Triton X-100 for 10 minutes, followed by blocking with 10% FBS for 1 hour at room temperature. Primary antibodies, including homemade anti-FOXG1-C-term (GP) and anti-FOXG1-N-term (Invitrogen #PA5-41493, Rabbit), were applied and left overnight at 4°C. After three PBT washes, secondary antibodies donkey-anti-guinea pig-647 and donkey-anti-rabbit-Cy3 were used and incubated for 1 hour. Following DAPI staining and mounting, the imaging was performed using a Nikon Eclipse Ti2 confocal microscope.

### Statistical analyses

The data were analyzed using GraphPad 10 (Prism, San Diego, CA). Details about the test statistics were provided in Supplementary Information. For histological analyses, unpaired t-test and Mann-Whitney test were used for two-group comparisons after checking normal distribution. Behavioral data were analyzed with two-way repeated-measures ANOVA or two-way ANOVA mixed-effects model. Statistical significance was represented on the graphs as: ^∗∗∗∗^p < 0.0001; ^∗∗∗^p < 0.001; ^∗∗^p < 0.01; ^∗^p < 0.05; ns, not significant. All bars and error bars were specified in the figure legends.

### Data availability

The RNA-seq raw data generated in this study have been deposited in the NCBI’s Gene Expression Omnibus database under the accession number GSE225399, and the Supplementary Information files and the Source Data file are provided with this paper.

## Acknowledgments

We are grateful to Hyeryeong Park, Bongjin Shin, Jingyan Huang, Medha K C, and Maria Root for their various contributions to this study. Also we greatly appreciate the efforts and support of the staff and veterinarians of the University at Buffalo LAF. This work was funded by grants from NINDS/NIH (NS111760 and NS100471 to S.-K.L. and NS118748 to S.-K.L. and J.W.L) and FOXG1 Research Foundation (to S.-K.L. and J.W.L) as well as the generous startup fund from University at Buffalo (to S.-K.L. and J.W.L). The authors have no conflicts of interest to disclose.

## Notes

### Competing Interest Statement

The authors have declared no competing interest.

