## Supplemental figures and information for "The patient-specific mouse model with *Foxg1* frameshift mutation provides insights into the pathophysiology of FOXG1 syndrome"

**Supplementary figures and information**

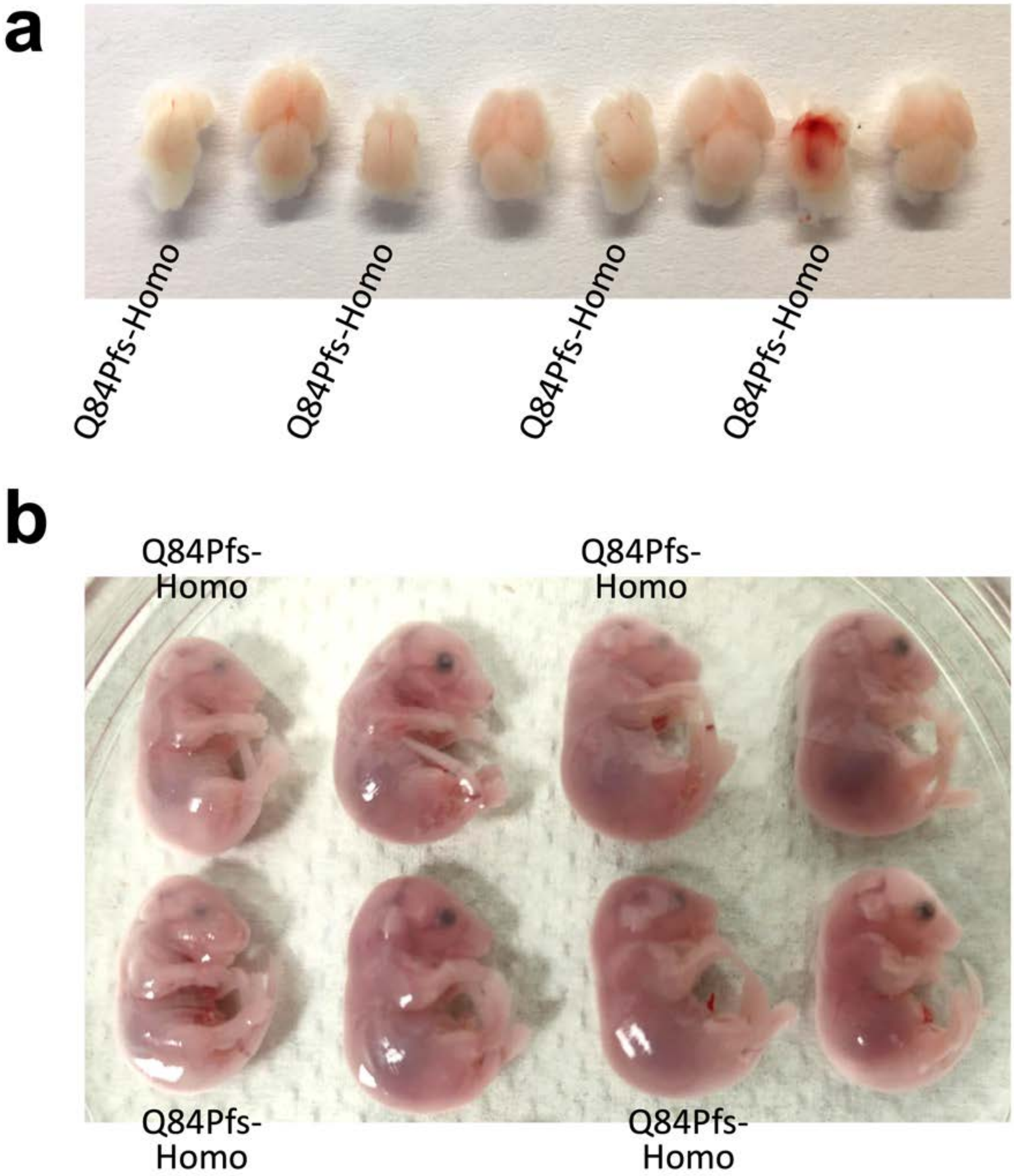


**Supplementary Fig. 1. The phenotypes of Q84Pfs-Homo mice. a**, Q84Pfs-Homo mice exhibited drastically reduced forebrain mass at E18.5. **b**, Q84Pfs-Homo mice exhibited craniofacial defects, such as underdeveloped frontonasal structures at E18.5.

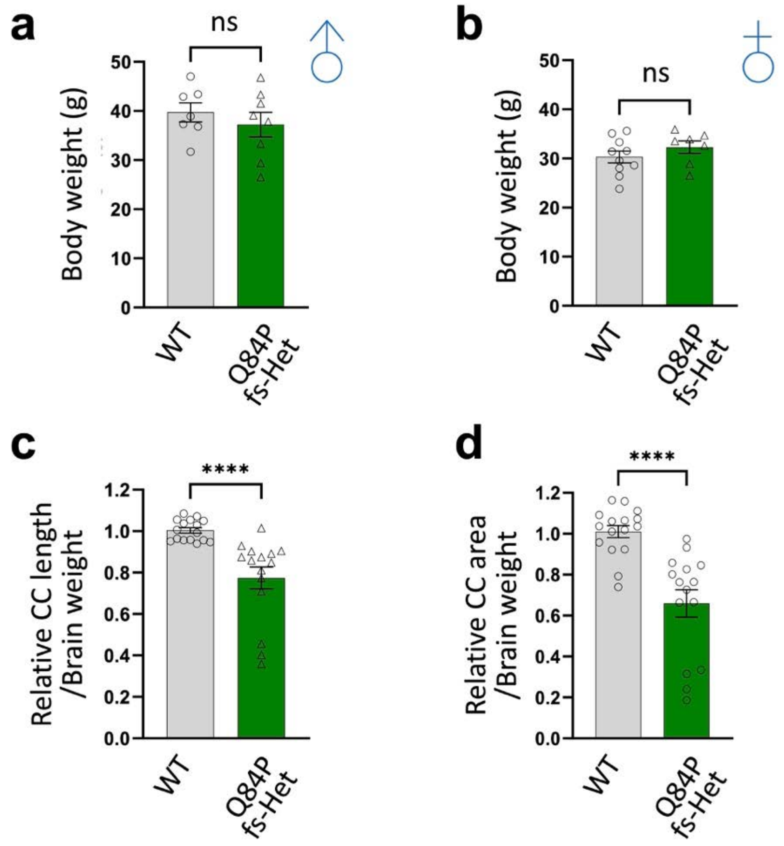


**Supplementary Fig. 2. Q84Pfs-Het adult mice exhibited corpus callosum reduction. a,b,** Quantification of body weight of adult male (**a**) and female (**b**) mice. **c,d,** Relative length (**c**) or area (**d**) of the corpus callosum (CC) normalized by brain weight. n=7 males, 9 females for WT, 8 males, and 7 females for Q84Pfs-Het mice. The error bars represent the standard deviation of the mean. ns, non-significant in unpaired two-tailed t test (a,b) and ****$p$<0.0001 in Mann-Whitney test (c,d).

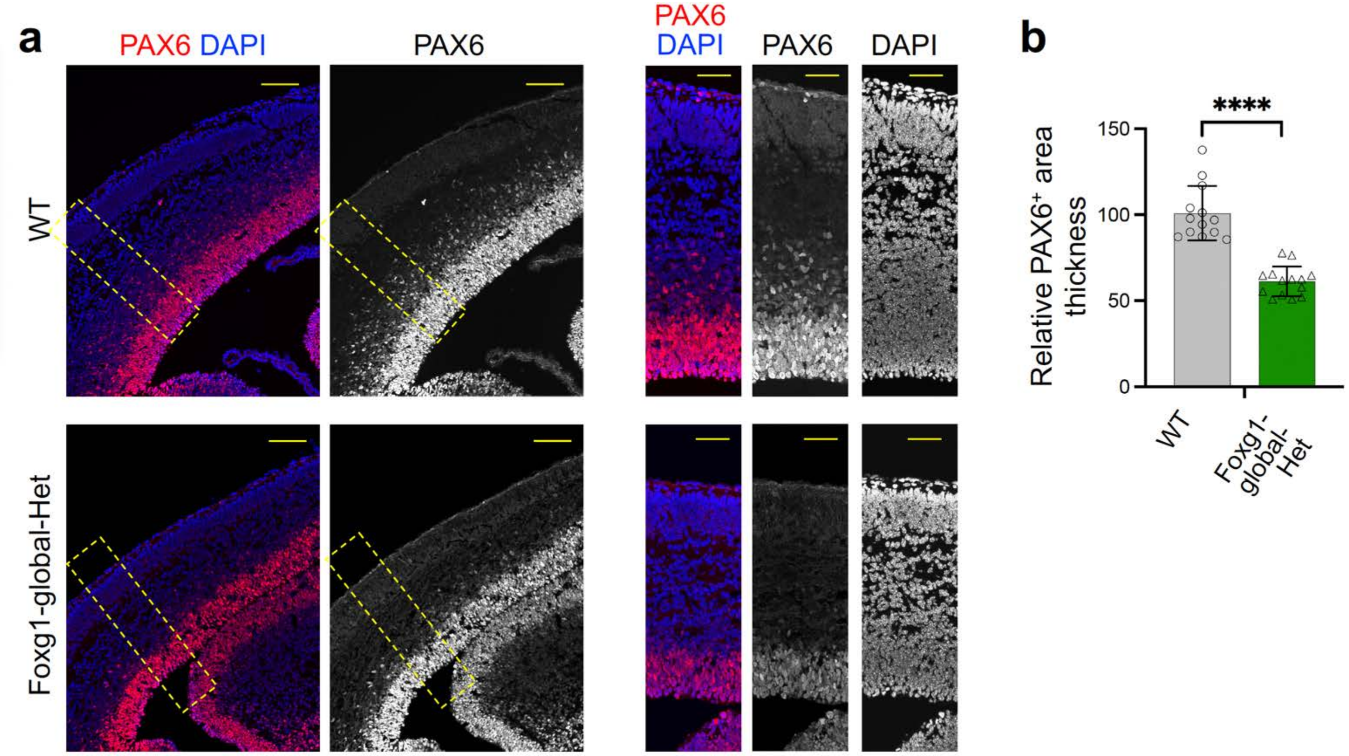


**Supplementary Fig. 3. Reduction of NPCs in global Foxg1-Het cortex.** The immunostaining analyses of global Foxg1-Het and WT cortices at E16 with the NPC marker Pax6. The thickness of Pax6$^{+}$ progenitor zones was measured in three independent areas per section (n = 4 mice/condition). Scale bars, 100μm (lower magnification images), or 50μm (higher magnification images). The error bars (**b**) represent the standard deviation of the mean. ****$p$<0.0001 in unpaired two-tailed t test. Only the representative images are shown.

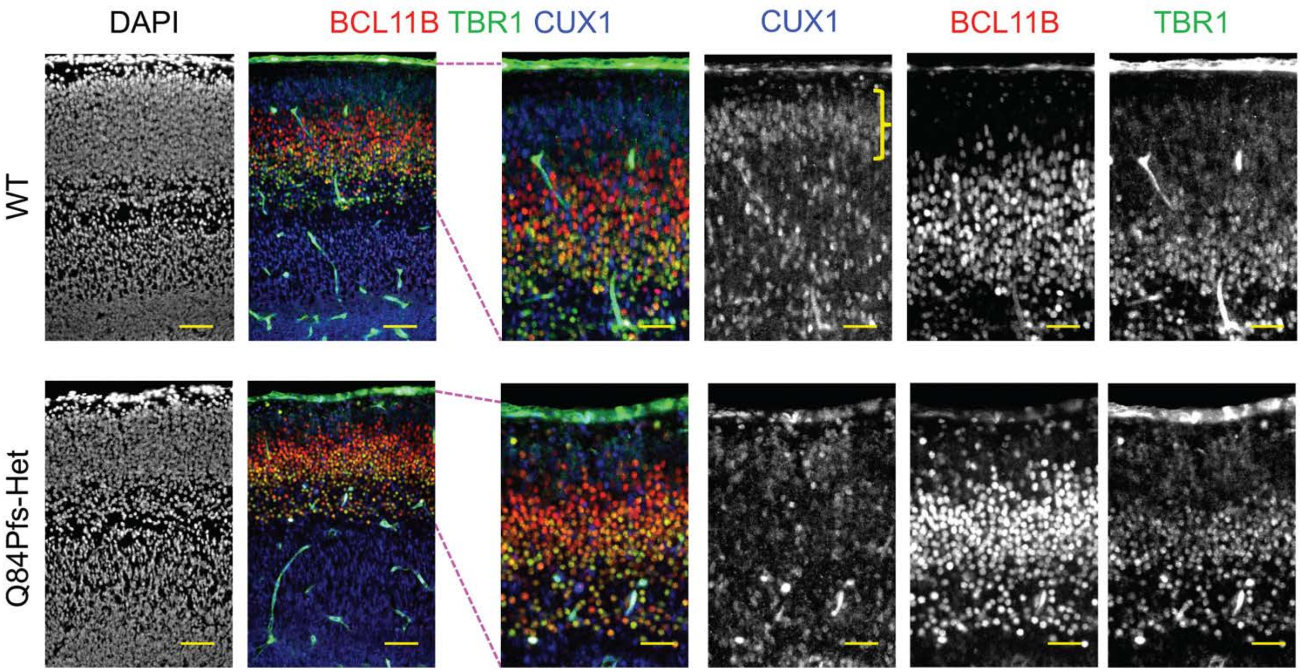


**Supplementary Fig. 4. The production of upper layer neurons was delayed in Q84Pfs-Het cortex.** The immunostaining analyses of Q84Pfs-Het and WT cortices at E16. The yellow bracket marks the $Cux1^{+}$ upper layer. Bcl11b and Tbr1 label deep layer neurons. Scale bars, 100µm (lower magnification images), or 20µm (higher magnification images). Only representative images are shown.

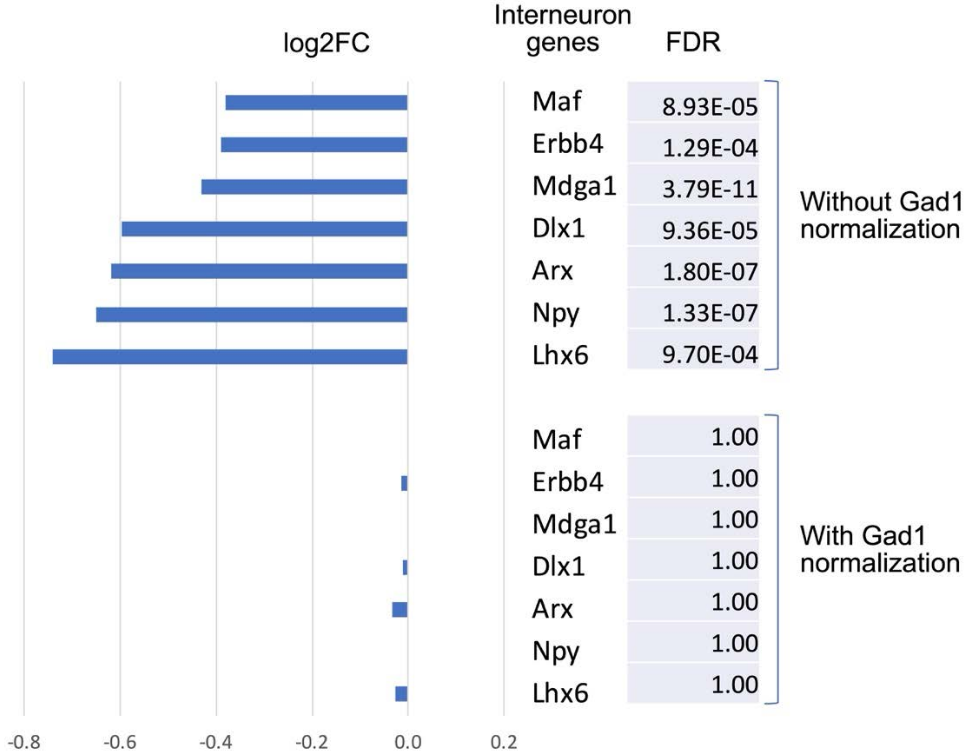


**Supplementary Fig. 5. The analysis of expression levels of cortical interneuron genes.** The analysis of RNA-seq data of P1 cortices. The X-axis represents the log2 fold changes (log2FC) of gene expression in Q84Pfs-Het compared to WT cortices. After normalization to Gad1 expression levels, the gene expression of interneurons was similar between Q84Pfs-Het and WT cortices. FDR stands for false discovery rate.

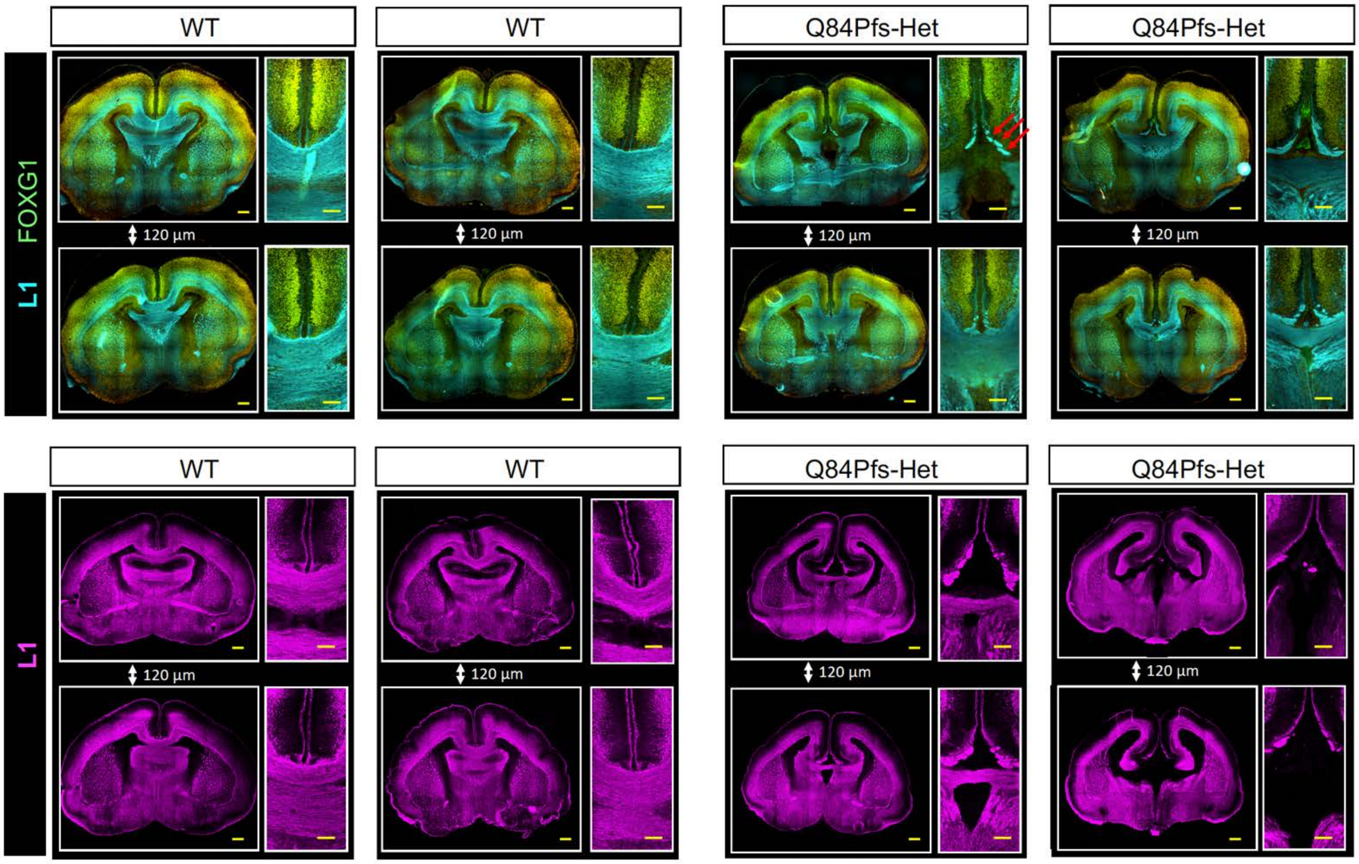


**Supplementary Fig. 6. Q84Pfs-Het cortex exhibited the prominent Probst bundles.** The immunostaining analyses of Q84Pfs-Het and WT brains with the axonal marker L1 at P0. The images of four WT and Q84Pfs-Het mice were displayed. For each mouse, a pair of brain images spaced 120µm apart was provided. Enlarged images of the midline region were placed adjacent to the whole brain coronal section images, highlighting the presence of the Probst bundle (aka, longitudinal callosal fascicles) exclusively in Q84Pfs-Het mice (indicated as red arrows only in the first Q84Pfs-Het brain) and its absence in WT mice. The images in the top and bottom rows originate from two different litters. Scale bars, 2mm (lower magnification images), or 1mm (higher magnification images).

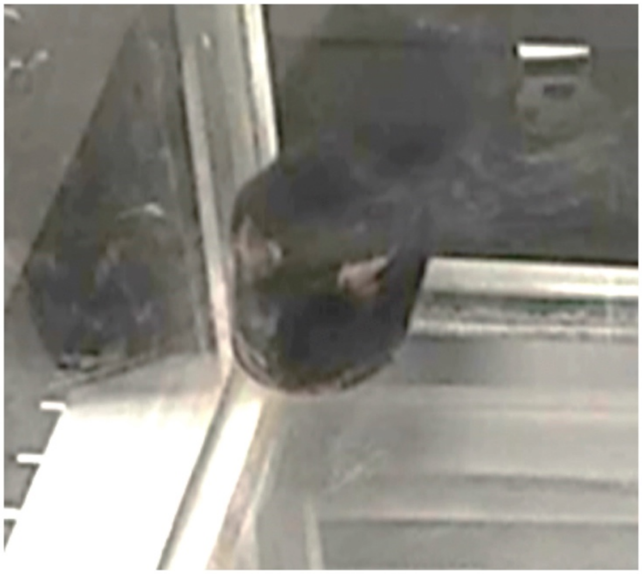

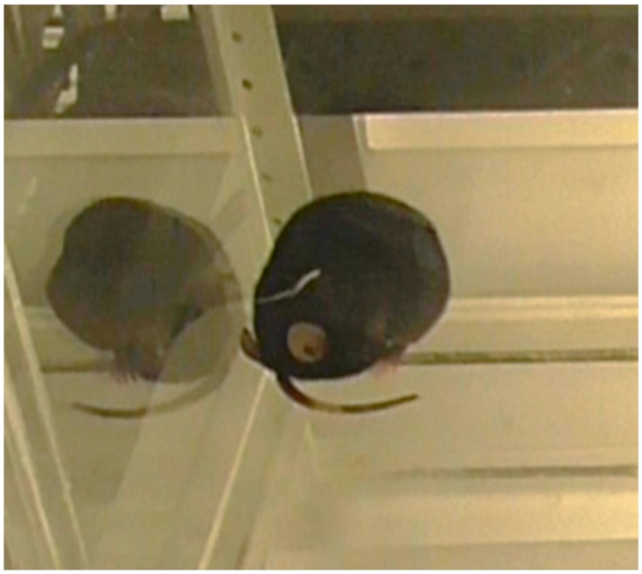

**Supplementary Fig. 7. Prolonged behavior arrest of Q84Pfs-Het mice.** Example photos of Q84Pfs-Het mice exhibiting prolonged behavioral arrest. Mice showing this behavior were always sitting in the corner of the cage and usually had their faces positioned outward towards the center of the cage (left). One mouse's body appeared quite hunched, and its head was positioned underneath its body for several minutes (right). Full videos are included in the supplementary information.

**Supplementary video 1:** Example of WT mouse behavior during the locomotor open field test (P60, female).

**Supplementary video 2:** Example of Q84Pfs-Het mouse exhibiting prolonged behavior arrest during the locomotor open field test (P60, female). This was the longest period of behavior arrest that was observed (approximately 14 minutes).

**Supplementary video 3:** Example of Q84Pfs-Het mouse exhibiting prolonged behavior arrest with abnormal body positioning during the locomotor open Field test (P90, female).

**Supplementary data, DEGs in Excel sheet**
222 differentially expressed genes (DEGs, 118 upregulated genes, and 104 downregulated genes) of Q84Pfs-Het cortex, relative to WT cortex, are listed.

**Statistics**
**Fig. 2**
g. Corpus callosum thickness: Mann-Whitney test
"AP-1: WT vs. Q84Pfs-Het" $p= 0.0079$
"AP-2: WT vs. Q84Pfs-Het" $p= 0.4127$
"AP-3: WT vs. Q84Pfs-Het" $p= 0.5556$
"AP-4: WT vs. Q84Pfs-Het" $p= 0.7302$
"AP-5: WT vs. Q84Pfs-Het" $p= 0.3333$

h. Dentate gyrus length: Unpaired two-tailed t test
$t= 10.57$, $df= 16$, $p< 0.0001$

i. Hippocampus area: Unpaired two-tailed t test
$t= 2.889$, $df= 16$, $p= 0.0107$

j. Body weight: Unpaired two-tailed t test
$t= 0.3307$, $df= 30$, $p= 0.7432$

k. Brain weight: Unpaired two-tailed t test
$t= 7.220$, $df= 30$, $p< 0.0001$

m. Corpus callosum A-P length: Mann-Whitney test
"WT vs. Q84Pfs-Het" $p< 0.0001$

n. Corpus callosum area: Mann-Whitney test
"WT vs. Q84Pfs-Het" $p< 0.0001$

**Fig. 3**
b. Mean intensity: Mann-Whitney test

"FOXG1-N-Ab: WT vs. Q84Pfs-Het" p= 0.0011
"FOXG1-C-Ab: WT vs. Q84Pfs-Het" p< 0.0001

**Fig. 4**
b. *In utero* electroporation GFP-positive cell ratio: Mann-Whitney test
"VZ: WT vs. Q84Pfs-Het" p= 0.5377
"IZ: WT vs. Q84Pfs-Het" p= 0.0027
"SP: WT vs. Q84Pfs-Het" p= 0.0650
"CP: WT vs. Q84Pfs-Het" p= 0.0045

d. GFP+ RGC's glial scaffold counting: Mann-Whitney test
"pCIG vs. Q86Pfs" p< 0.0001

f. Double positive BrdU+ GFP+ cell ratio: Mann-Whitney test
"pCIG vs. Q86Pfs" p< 0.0001

**Fig. 6**
b. Cortex thickness: Unpaired two-tailed t test
t= 4.340, df= 52, p< 0.0001

c. PAX6+ area thickness: Unpaired two-tailed t test
t= 6.571, df= 52, p< 0.0001

e. pHH3+ cell counting: Unpaired two-tailed t test
t= 2.968, df= 4, p= 0.0412

**Fig. 7**
b. Marker-positive cell counting at E16: Unpaired two-tailed t test
"BCL11B-positive: WT vs. Q84Pfs-Het" t= 2.325, df= 49, p= 0.0243
"CUX1-positive: WT vs. Q84Pfs-Het" t= 6.558, df= 49, p< 0.0001

d. Layer thickness at P1: Unpaired two-tailed t test
"Deep layer: WT vs. Q84Pfs-Het" t= 0.2056, df= 43, p= 0.8381
"Upper layer: WT vs. Q84Pfs-Het" t= 8.174, df= 43, p< 0.0001

e. Marker-positive cell counting at P1: Unpaired two-tailed t test
"TBR1-positive: WT vs. Q84Pfs-Het" t= 2.924, df= 15, p= 0.0105
"BCL11B-positive: WT vs. Q84Pfs-Het" t= 2.834, df= 15, p= 0.0126
"CUX1-positive: WT vs. Q84Pfs-Het" t= 3.552, df= 15, p= 0.0029

g. Relative Cortex thickness: Unpaired two-tailed t test
t= 8.319, df= 40, p< 0.0001

h. Relative upper layer thickness: Unpaired two-tailed t test
t= 5.219, df= 14, p= 0.0001

i. Relative deep layer thickness: Unpaired two-tailed t test
t= 1.283, df= 22, p= 0.2130

k. DLX1+ cell counting: Unpaired two-tailed t test

t= 4.844, df= 6, p= 0.0029

**Fig. 8**
c. Vertical L1+ axon bundles/area at E16: Unpaired two-tailed t test
t= 6.041, df= 6, p= 0.0009

d. Vertical L1+ axon bundles/area at P1: Unpaired two-tailed t test
t= 7.471, df= 6, p= 0.0003

**Fig. 9**
b. OLIG2+ cell counting: Unpaired two-tailed t test
t= 6.152, df= 37, p< 0.0001

d. MBP+ area ratio: Unpaired two-tailed t test
t= 9.083, df= 35, p< 0.0001

e. Relative MBP intensity: Unpaired two-tailed t test
t= 6.482, df= 22, p< 0.0001

**Fig. 10**
c. Body weight: Two-way repeated measures ANOVA
<Male>
Interaction: F(2,38)= 3.259, p= 0.0494
Age (P30, P60, or P90): F(2,38)= 685.4, p< 0.0001
Genotype (WT or Q84Pfs-Het): F(1,19)= 0.009348, p= 0.9240
Post-hoc (Sidak)
"P30: WT vs. Q84Pfs-Het" p= 0.4538, Mean diff.= 0.8806, 95.00% CI of diff.= -0.7247 to 2.486
"P60: WT vs. Q84Pfs-Het" p= 0.3158, Mean diff.= -1.033, 95.00% CI of diff.= -2.639 to 0.5719
"P90: WT vs. Q84Pfs-Het" p= 0.9595, Mean diff.= 0.2917, 95.00% CI of diff.= -1.314 to 1.897
<Female>
Interaction: F(2,34)= 0.8790, p= 0.4244
Age (P30, P60, or P90): F(1.281,21.78)= 137.9, p< 0.0001
Genotype (WT or Q84Pfs-Het): F(1,17)= 2.256, p= 0.1514
Post-hoc (Sidak)
"P30: WT vs. Q84Pfs-Het" p= 0.9843, Mean diff.= 0.3367, 95.00% CI of diff.= -2.562 to 3.235
"P60: WT vs. Q84Pfs-Het" p= 0.2616, Mean diff.= 1.409, 95.00% CI of diff.= -0.7821 to 3.600
"P90: WT vs. Q84Pfs-Het" p= 0.3758, Mean diff.= 1.906, 95.00% CI of diff.= -1.490 to 5.301

d. Wire hanging: Two-way ANOVA
Interaction: F(2,76)=3.710, p=0.0290
Age (P30, P60, or P90): F(1.684,64.00)= 2.797, p= 0.0772
Genotype (WT or Q84Pfs-Het): F(1,40)= 14.16, p= 0.0005
Post-hoc (Sidak)
"P30: WT vs. Q84Pfs-Het" p= 0.3674, Mean diff.= 125.3, 95.00% CI of diff.= -82.93 to 333.6
"P60: WT vs. Q84Pfs-Het" p= 0.0032, Mean diff.= 309.5, 95.00% CI of diff.= 92.78 to 526.3
"P90: WT vs. Q84Pfs-Het" p= 0.0008, Mean diff.= 310.3, 95.00% CI of diff.= 123.0 to 497.6

e. Travel distance in Open field test: Two-way repeated measures ANOVA
For P30,
Interaction: F(5,180)= 8.418, p< 0.0001

Time (min): F(3.350,120.6)= 35.36, p< 0.0001
Genotype (WT or Q84Pfs-Het): F(1,36)= 0.01682, p= 0.8975
Post-hoc (Sidak)
“10 min: WT vs. Q84Pfs-Het” p= 0.1764, Mean diff.= -310.7, 95.00% CI of diff.= -699.8 to 78.45
“20 min: WT vs. Q84Pfs-Het” p= 0.2451, Mean diff.= -241.2, 95.00% CI of diff.= -565.9 to 83.50
“30 min: WT vs. Q84Pfs-Het” p= 0.9975, Mean diff.= -48.40, 95.00% CI of diff.= -328.4 to 231.6
“40 min: WT vs. Q84Pfs-Het” p= 0.9639, Mean diff.= 90.37, 95.00% CI of diff.= -221.5 to 402.2
“50 min: WT vs. Q84Pfs-Het” p= 0.6118, Mean diff.= 186.3, 95.00% CI of diff.= -167.1 to 539.8
“60 min: WT vs. Q84Pfs-Het” p= 0.1409, Mean diff.= 253.4, 95.00% CI of diff.= -48.30 to 555.0

For P60,
Interaction: F(5,180)= 3.649, p= 0.0036
Time (min): F(3.315,119.3)= 92.91, p< 0.0001
Genotype (WT or Q84Pfs-Het): F(1,36)= 30.01, p< 0.0001
Post-hoc (Sidak)
“10 min: WT vs. Q84Pfs-Het” p= 0.2737, Mean diff.= 258.4, 95.00% CI of diff.= -100.5 to 617.3
“20 min: WT vs. Q84Pfs-Het” p= 0.0001, Mean diff.= 416.3, 95.00% CI of diff.= 183.6 to 649.0
“30 min: WT vs. Q84Pfs-Het” p< 0.0001, Mean diff.= 562.4, 95.00% CI of diff.= 254.4 to 870.5
“40 min: WT vs. Q84Pfs-Het” p= 0.0010, Mean diff.= 522.0, 95.00% CI of diff.= 179.2 to 864.9
“50 min: WT vs. Q84Pfs-Het” p= 0.0003, Mean diff.= 555.0, 95.00% CI of diff.= 221.1 to 889.0
“60 min: WT vs. Q84Pfs-Het” p <0.0001, Mean diff.= 651.3, 95.00% CI of diff.= 343.5 to 959.1

For P90,
Interaction: F(5,185)= 2.866, P= 0.0162
Time (min): F(1.958,72.44)= 56.44, p< 0.0001
Genotype (WT or Q84Pfs-Het): F(1,37)= 12.35, P= 0.0012
Post-hoc (Sidak)
“10 min: WT vs. Q84Pfs-Het” p= 0.9821, Mean diff.= 135.0, 95.00% CI of diff.= -410.2 to 680.2
“20 min: WT vs. Q84Pfs-Het” p= 0.6561, Mean diff.= 265.9, 95.00% CI of diff.= -264.0 to 795.9
“30 min: WT vs. Q84Pfs-Het” p= 0.0631, Mean diff.= 353.8, 95.00% CI of diff.= -12.70 to 720.2
“40 min: WT vs. Q84Pfs-Het” p= 0.0027, Mean diff.= 548.5, 95.00% CI of diff.= 158.0 to 938.9
“50 min: WT vs. Q84Pfs-Het” p= 0.0239, Mean diff.= 505.4, 95.00% CI of diff.= 48.41 to 962.3
“60 min: WT vs. Q84Pfs-Het” p= 0.0002, Mean diff.= 631.0, 95.00% CI of diff.= 259.1 to 1003

g. Center time in Open field test: Two-way ANOVA
Interaction: F(2,72)= 2.333, p= 0.1043
Age (P30, P60, or P90): F(1.717,61.82)= 7.072, p= 0.0028
Genotype (WT or Q84Pfs-Het): F(1,37)= 20.17, p< 0.0001
Post-hoc (Sidak)
“P30: WT vs. Q84Pfs-Het” p= 0.0681, Mean diff.= 141.1, 95.00% CI of diff.= -7.960 to 290.2
“P60: WT vs. Q84Pfs-Het” p= 0.0007, Mean diff.= 253.2, 95.00% CI of diff.= 101.7 to 404.7
“P90: WT vs. Q84Pfs-Het” p< 0.0001, Mean diff.= 235.6, 95.00% CI of diff.= 120.5 to 350.7

h. Average duration of grooming in Open field test: Mann-Whitney test
“P30: WT vs. Q84Pfs-Het” p= 0.003179
“P60: WT vs. Q84Pfs-Het” p= 0.000063
“P90: WT vs. Q84Pfs-Het” p= 0.000194

i. Marble burying test: Two-way ANOVA
Interaction: F(2,76)= 48.99, p< 0.0001

Age (P30, P60, or P90): F(1.947,73.98)= 83.71, p< 0.0001
Genotype (WT or Q84Pfs-Het): F(1,40)= 12.39, p= 0.0001
Post-hoc (Sidak)
"P30: WT vs. Q84Pfs-Het" p= 0.0896, Mean diff.= 2.823, 95.00% CI of diff.= -0.3338 to 5.979
"P60: WT vs. Q84Pfs-Het" p< 0.0001, Mean diff.= -6.880, 95.00% CI of diff.= -9.802 to -3.957
"P90: WT vs. Q84Pfs-Het" p< 0.0001, Mean diff.= -7.038, 95.00% CI of diff.= -10.05 to -4.029

**Supplementary Fig. 2.**
a. Body weight of adult male: Unpaired two-tailed t test
t= 0.7807, df= 13, p= 0.4490

b. Body weight of adult female: Unpaired two-tailed t test
t= 1.100, df= 15, p= 0.2885

c. Relative corpus callosum length normalized by brain weight: Mann-Whitney test
"WT vs. Q84Pfs-Het" p< 0.0001

d. Relative corpus callosum area normalized by brain weight: Mann-Whitney test
"WT vs. Q84Pfs-Het" p< 0.0001

**Supplementary Fig. 3.**
a. Relative PAX6 area thickness: Unpaired two-tailed t test
t= 8.174, df= 25, p< 0.0001
